# Differential regulation of the immune system in a brain-liver-fats organ network during short term fasting

**DOI:** 10.1101/2020.04.05.026351

**Authors:** Susie S.Y. Huang, Melanie Makhlouf, Eman H. AbouMoussa, Mayra L. Ruiz Tejada Segura, Lisa S. Mathew, Kun Wang, Man C. Leung, Damien Chaussabel, Darren W. Logan, Antonio Scialdone, Mathieu Garand, Luis R. Saraiva

## Abstract

Different fasting regimens are known to promote health, mitigate chronic immunological disorders, and improve age-related pathophysiological parameters in animals and humans. Indeed, several clinical trials are currently ongoing using fasting as a potential therapy for a wide range of conditions. Fasting alters metabolism by acting as a reset for energy homeostasis. However, the molecular mechanisms underlying the beneficial effects of short-term fasting (STF) are still not well understood, particularly at the systems or multi-organ level. Here, we investigated the dynamic gene expression patterns associated with six periods of STF in nine different mouse organs. We cataloged the transcriptional dynamics within and between organs during STF and discovered differential temporal effects of STF among organs. Using gene ontology enrichment analysis, we identified an organ network sharing 37 common biological pathways perturbed by STF. This network incorporates the brain, liver, interscapular brown adipose tissue, and posterior-subcutaneous white adipose tissue, hence we named it the brain-liver-fats organ network. Using Reactome pathways analysis, we identified the immune system, dominated by T cell regulation processes, as a central and prominent target of systemic modulations during STF in this organ network. The changes we identified in specific immune components point to the priming of adaptive immunity and parallel the fine-tuning of innate immune signaling. Our study provides a comprehensive multi-organ transcriptomic profiling of mice subjected to multiple periods of STF, and adds new insights into the molecular modulators involved in the systemic immuno-transcriptomic changes that occur during short-term energy loss.

## INTRODUCTION

Dietary restriction refers to an intervention that can range from chronic but minor reduction in calorie intake to periods of repeated cycles of short-term fasting (STF) and, in humans, those lasting more than 48h [1]. These various forms of reduction in food consumption have many beneficial effects on health, including weight reduction, amelioration of auto-immune diseases, and increased lifespan [2,3]. In line with this, a triad of recent studies demonstrated the ability of different forms of fasting on changing the level and functions of the various immune cell types [4–6]. Recently, the interest and applicability of fasting to treat various conditions are as such that they have generated momentum in clinical trials [7–9].

The advent of high-throughput omics facilitated the elucidation of some of the cellular and molecular mechanisms underlying the beneficial effects of fasting. Nonetheless, the majority of these studies focused on single organ response (i.e., the liver) and, less often, involved two or three organs [10–16]. The adaptation to energy deprivation, however, requires a multi-organ integration of metabolic modulations to protect the organism from an irreversible loss of resources [17]. In mice, disturbance of normal eating patterns alters metabolism systemically [18]. A recent study described the interorgan coordination of adaptive responses to various periods of fasting (0-72 hours) in mice; however, the transcriptomic changes characterized in the four tissues applied mainly to states of starvation and are limited by the depth offered by microarrays [19]. Another recent multi-omic approach also presented similar limitations [20]. Currently, a systemic or multi-organ overview of the in-depth molecular changes (i.e., transcriptomics) mediated by fasting is lacking.

In this study, we profiled the transcriptomic responses to periods of STF (0-22 hours) in nine different mouse organs. Using a combination of data-driven and semantic similarity clustering approaches, we discovered the presence of a brain-liver-fats organ network, conserved in their enriched biological processes perturbed by STF. The organ network recapitulated numerous reported physiological/molecular changes associated with various forms of fasting. In all, we provide a multi-organ consensus of known and novel molecular mediators of systemic effects of short-term energy loss in mice.

## RESULTS

### Transcriptome profiling across multiple organs in the fasted mice

To investigate the global gene expression dynamics associated with STF, we used mRNA-seq to profile the transcriptomes of nine organs obtained from mice subjected to six different STF duration (0, 2, 8, 12, 18 and 22 hours of fasting; n=3 per time point; Fig. 1a). The nine organs profiled were: olfactory bulb (OB), brain (BRN, which includes the telencephalon and diencephalon), cerebellum (CBL), brainstem (BST, which consists of the mesencephalon, pons, and myelencephalon), stomach (STM), liver (LIV), interscapular brown adipose tissue (iBAT), perigonadal white adipose tissue (pgWAT), and posterior-subcutaneous white adipose tissue (psWAT). After quality control (see Methods), we retained 157 out of 162 samples (97%) for downstream analyses. After applying a cutoff of >10 normalized counts in at least three samples, we found that 13,129-15,012 genes were expressed per organ (Additional file 1: Fig. S1; Additional file 2: Table S1). Next, we performed a principal component analysis on the genes expressed across all samples (9420) and found segregation primarily according to organ type and into four different clusters: one composed by all nervous system samples (OB, BRN, CBL, BST), one by all adipose tissues (iBAT, pgWAT, psWAT), and the two remaining clusters composed by STM and LIV, respectively (Fig. 1b). Principal component analyses made on all samples from each organ also did not yield clear separation by fasting time (data now shown). A hierarchical clustering analysis of the union of the top 100 most abundant genes expressed across all samples also resulted in similar sample clustering (Fig. 1c). Together these results indicate that organ clustering is primarily driven by anatomical similarity, consistent with previous findings [21–23].

**Figure 1.**
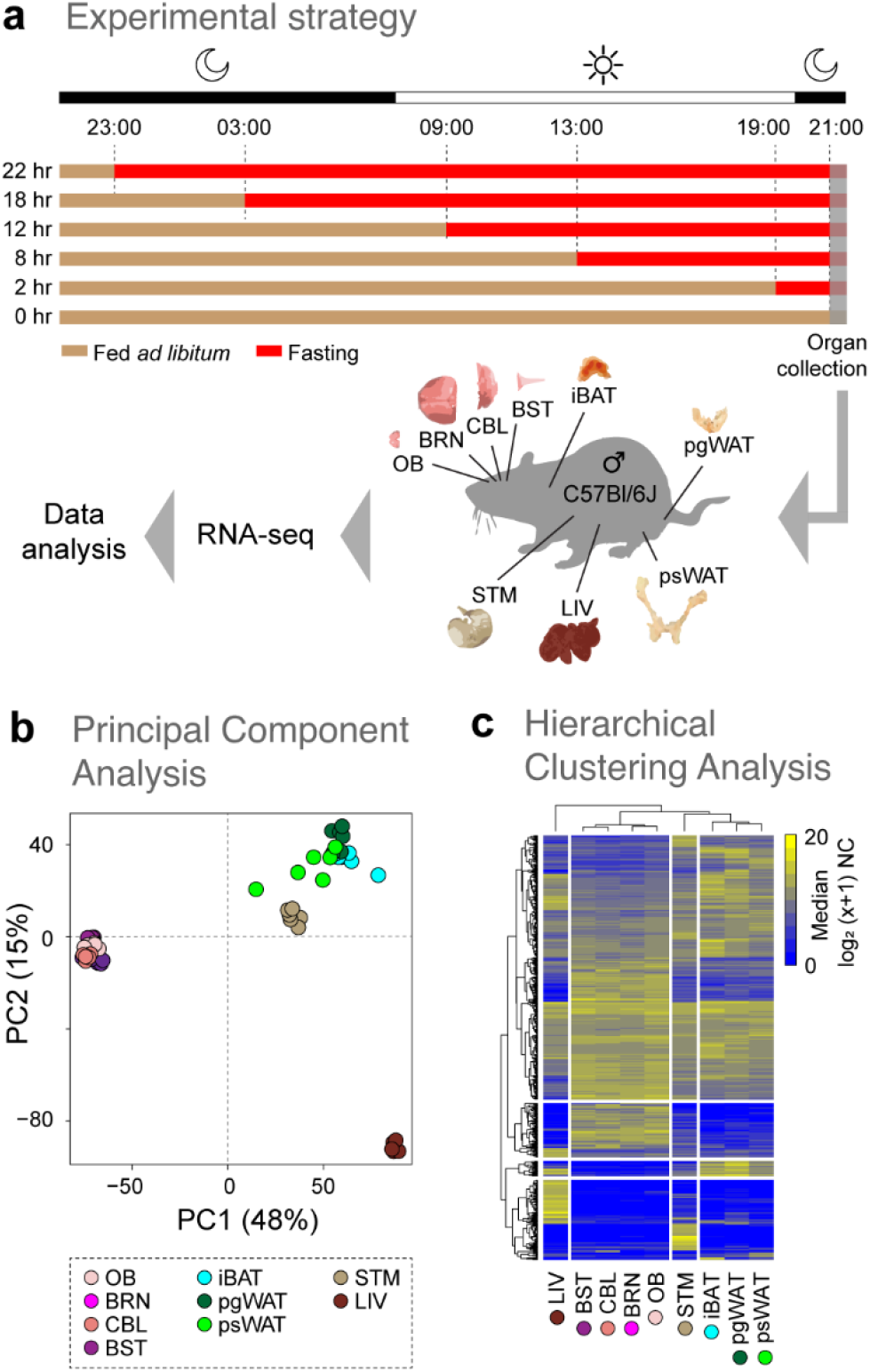
Transcriptomic profiling of multiple organs in the fasted mice. **a)** Schematic view of the experimental design for the six different short term fasting (STF) durations are indicated by the red bars (0, 2, 8, 12, 18, and 22 hours). The fed ad libitum condition is shown by the beige bars. The time of organ collection from male C57BL/6J mice (n=3 per time point) is represented by the grey-shaded area. Periods of light-on and light-off are represented by the white and black bars, respectively. **b)** Principal component (PC) analysis of the 9420 expressed genes in all samples. Each dot represents the gene expression profile of an organ (indicated by the colour) at a specific time point. Percentages of the variance explained by the PCs are indicated in parentheses. **c)** Hierarchical clustering analysis of the union on the top 100 highly expressed genes (rows) among all organs (columns). Median mRNA expression levels are represented on a log_2_ (x+1) scale of normalized counts (NC) (0 - not expressed; 20 - highly expressed). Organ abbreviations: OB - olfactory bulb, BRN - brain (which includes the telencephalon and diencephalon), CBL - cerebellum, BST - brainstem (which includes the mesencephalon, pons, and myelencephalon), STM - stomach, LIV - liver, iBAT - interscapular brown adipose tissue, pgWAT - perigonadal white adipose tissue, and psWAT - posterior-subcutaneous white adipose tissue.

### Data-driven clustering shows differential temporal effects of STF among organs

To examine the temporal impact of STF for each organ, we employed a data-driven approach aimed at identifying time-dependent fasting clusters. To do so, we defined the highly variable genes (HVGs) within each organ and calculated the optimal number of clusters into which all samples segregated based on organ-specific HVGs (see Methods and Additional file 2: Fig. S1). We observed that for most organs (BRN, CBL, BST, STM, LIV and iBAT), samples grouped in three clusters, which we designated as Phase I, II, and III (Fig. 2a, Additional file 3: Table S2). Samples from OB and psWAT grouped only in two clusters (Phases I and III), which seemingly cycle throughout the 22 hours. This cyclical transcriptomic pattern is reminiscent and consistent with other cyclical physiological or metabolic responses occurring in these organs during fasting [24–26]. In contrast with these results, we failed to identify robust clusters using the samples from pgWAT (thus excluding it from our downstream analyses). While intriguing, the fact that pgWAT and psWAT show differential gene expression dynamics to STF is expected, as different adipose depots are functionally distinct and can display differential transcriptomic responses to fasting [27,28].

**Figure 2.**
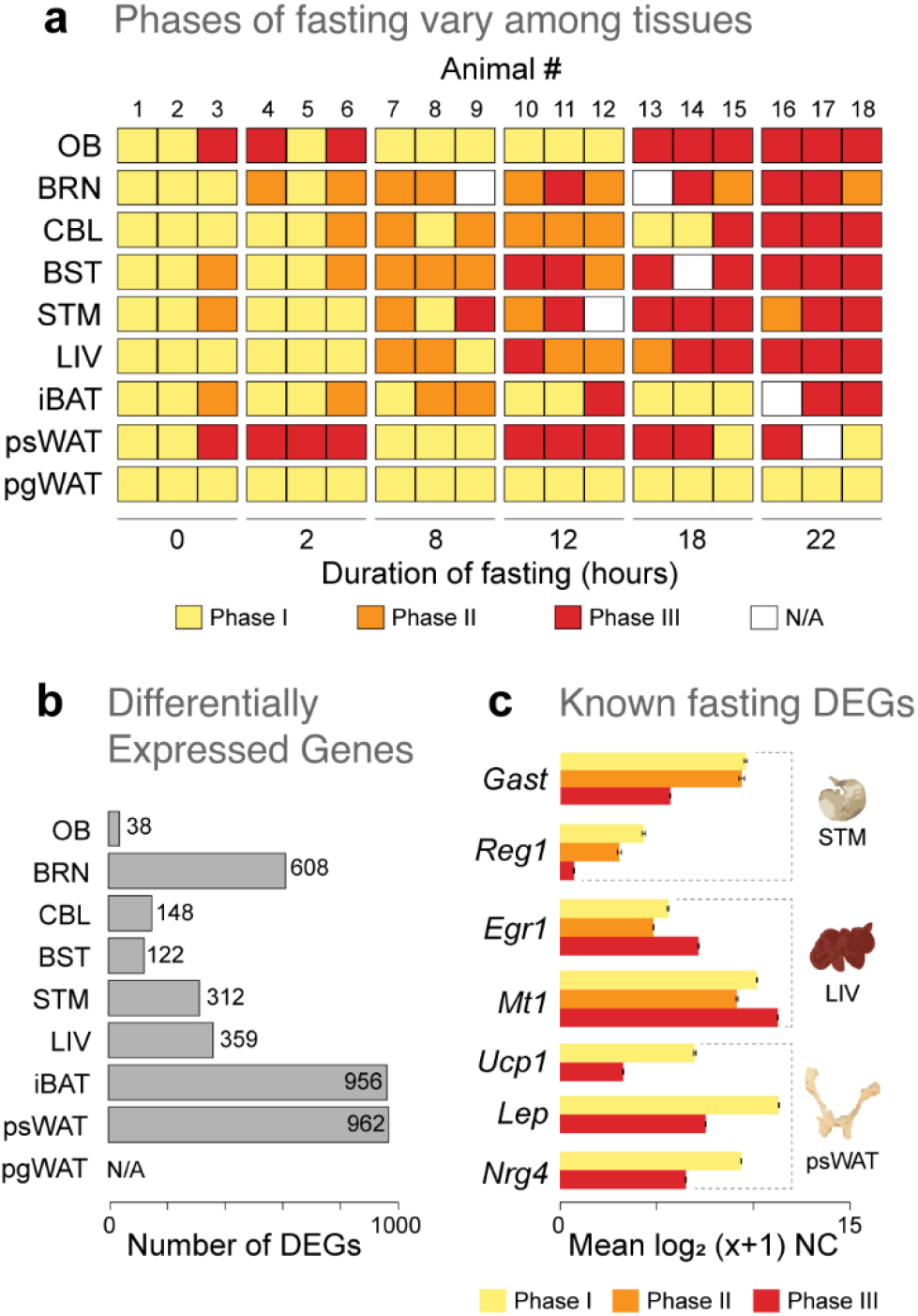
Gene expression-driven sample clustering shows differential temporal effects of STF among organs. a) Fasting Phase-call results from the data-driven clustering approach for each animal (columns) and organs (rows): Phase I (yellow), Phase II (orange), Phase III (red). N/A (white) represents excluded samples that did not meet the QA/QC criteria (see Additional file 3: Table S2). Experimental durations of fasting are as indicated. b) Number of differentially expressed genes (DEGs) from the all pairwise fasting Phase comparisons in all organs. Since samples from pgWAT did not split into fasting Phases, no DEGs for the organ were available (N/A). c) Expression values across the three Phases of select DEGs that have been previously reported to be modulated by fasting in selected organs. Mean mRNA expression levels are represented on a log_2_ (x+1) scale of normalized counts (NC) ± SEM (for replicate number, see Additional file 3: Table S2). Organ acronyms are as in Fig. 1.

To gain further insight into the transcriptomic changes associated with fasting, we performed a differential expression analysis for all pairwise Phase comparisons for each organ. The numbers of differentially expressed genes (DEGs; FDR<0.05, log2FC>1) among the organs ranged from 38 in the OB to 3826 in psWAT (Additional file 5: Table S4). To keep our analysis limited on the most impacted genes by STF, we focused our downstream analyses only on the top and bottom 12.5% of the log2FC ranked genes for organs yielding >500 DEGs (i.e., BRN, STM, iBAT, and psWAT). Hereafter, DEGs refer to these filtered DEGs.

The total number of DEGs identified for each organ varied greatly (Fig. 2b). While OB and BST displayed the lowest quantity of DEGs (38 and 122 respectively), the adipose tissues psWAT and iBAT showed the highest number of DEGs (962 and 956 respectively; Additional file 4: Table S3). This analysis yielded multiple sets of both known and newly identified genes that are affected by STF. For instance, between Phases I and III we found decreasing expression levels of *Nrg4, Lep* and *Ucp1* in psWAT, decreasing expression levels of Reg1 in STM, and increased expression of *Egr1, Fam107a, Map3k6*, and *Mt1* in the LIV (Fig. 2c; Additional file 5: Table S4), consistent with previous STF studies in rodents [11,20,29–32]. In sum, by allowing the similarity of HVGs among the samples to drive their clustering in an organ-specific manner, we were able to detect differential temporal effects of STF on the transcriptional dynamics across multiple organs.

### Organ-specific modulations by STF

To get insight into the functional roles of the hundreds of DEGs identified above, we combined multiple strategies to perform gene ontology (GO) enrichment analyses (see Methods). The biological pathway analysis returned significant results for psWAT, iBAT, LIV, BST, STM, and BRN, but not for CBL and OB (Fig. 3a). STF appears to induce changes in unique biological processes among the organs, with only two overlapping GO terms between LIV-psWAT and iBAT-psWAT, respectively (Fig. 3a, left panel). It is to note that the relatively small numbers of GO terms shown reflect the stringent thresholds and semantic reduction applied (see Methods); thus, only the most relevant and non-redundant terms are kept.

**Figure 3.**
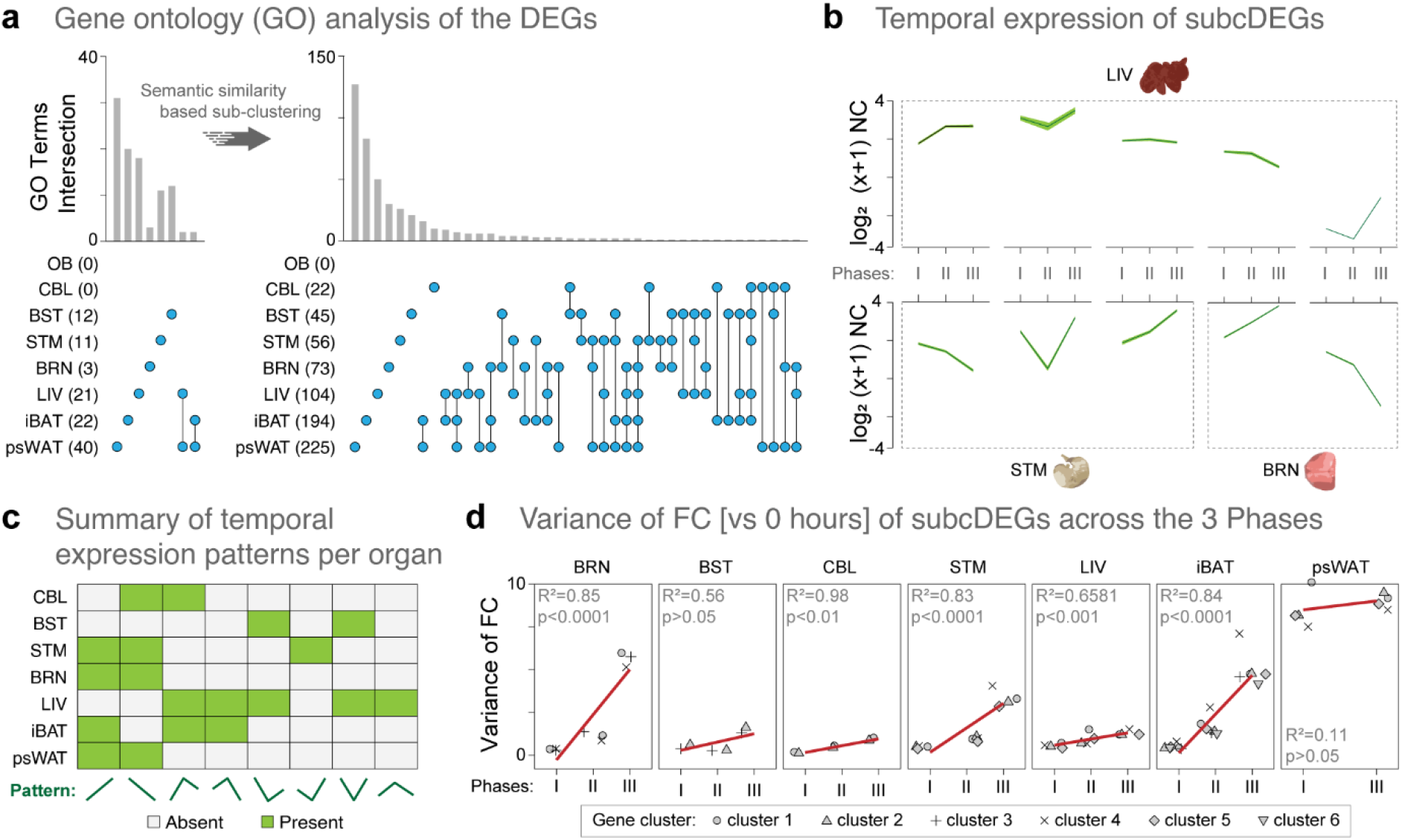
Functional re-clustering of DEGs increased the identification of the biological perturbations and the unique gene expression patterns in organs of the fasted mice. **a)** The number of gene ontology (GO) terms shared among organs increased after applying a semantic similarity based sub-clustering algorithm to the DEGs. GO analysis for the original DEGs, and for the subcDEGs are represented in the left and right panels, respectively. The number and nature of the GO intersections are denoted by the vertical bar graph and the line-dots, respectively. Number of GOs enriched from DEGs and subcDEGs are noted in between parenthesis in front of the organ acronym. **b)** The unique temporal expression patterns of the genes (as log_2_(x+1) NC) comprised in the subcDEGs from three selected organs (LIV, STM, BRN). mRNA expression levels are represented on a log_2_ (x+1) scale of mean normalized counts (NC) (dark green line) ± SD (light green shadow). **c)** Summary of the unique temporal expression patterns of the genes (as log_2_(x+1) NC) comprised in the subcDEGs in the seven organs. A total of eight different temporal expression patterns are present (green). **d)** Mean variance of the log_2_ fold change (FC) vs. timepoint 0 hour of the genes in all the subcDEGs across the three Phases. Linear regression analysis (red line) was performed for each organ, and the R^2^ and p-values are depicted in each panel. Up to six gene clusters, i.e. subcDEGs, have been identified among the seven organs; the cluster number is indicated at the bottom of the panel. Organ acronyms are as in Fig. 1.

GO semantic similarity provides the basis for functional comparison of gene products and are widely used in bioinformatics applications such as in cluster analysis of genes [33,34]. To improve the sensitivity and coverage of our GO enrichment analyses, we sub-clustered the organ-specific DEGs lists based on the semantic similarity of their associated GO terms (see Methods); hereafter, referred to as the sub-clustered DEGs (subcDEGs). We then performed GO enrichment for each of the multiple subcDEGs groups created for each organ; subcDEGs that did not yield enrichment of GO were excluded from downstream analysis. This new approach yielded a 5- to 24-fold increase in the enriched GO terms for all organs except OB, for which we still failed to identify enriched pathways, thus excluding it from further analysis (Fig. 3a, right panel; Additional file 6: Table S5). Notably, 22 GO terms were returned for CBL, which previously had no significant enrichments, and for BRN, the annotation coverage increased 24-fold, from 3 to 73 GO terms. To validate this new approach, we used a semantic network to visualize the GO terms enriched in LIV, which is the most well-studied tissue in the context of fasting (Additional file 1: Fig. S2). Among those, we found several GO terms consistent with previous fasting studies, including carboxylic acid biosynthetic process, lipid homeostasis, xenobiotic metabolism, regulation of protein stability, and regulation of protein localization to membrane [13,16,35]. Interestingly, we also identified other novel pathways enriched by STF, such as wound healing, sensory perception of pain and positive regulation of vasculature development, thus highlighting the discovery potential of this approach. In sum, by functionally re-clustering the DEGs, we have substantially increased both the organ-specific GO annotation coverage and the probability of discovering novel pathways associated with STF.

We then asked how the genes in each of the subcDEGs that provided GO enrichment varied over STF duration. To do so, we extracted and summarized the unique expression patterns of the collective subcDEGs against STF phases in each organ (Fig. 3b and c). In general, the non-brain organs exhibited a relatively higher number of distinct cluster expression patterns, suggesting comparatively higher dynamics in their gene expression response to STF. As expected, only two patterns were found in psWAT (explained by only two Phases). To further quantify these gene expression temporal perturbations, we calculated the variance of the mean fold-change versus timepoint 0 hour of the genes in all the subcDEGs across the three Phases and performed regression analysis (Fig. 3d). We found positive correlations of fold change variance with STF in most organs, except for BST and psWAT, where the relationships are non-significant. Interestingly, STM, BRN, and iBAT exhibited a noticeably higher degree of fold change variances with STF (i.e., the slope of the regression) compared with those of psWAT, CBL, and LIV, suggesting differential regulation of their transcriptional patterns as STF progressed. Together, these results provide a second line of evidence that different organs show different dynamics in their transcriptional programs to STF. Moreover, these data suggest that organs from the nervous and digestive/adipose systems use different strategies to cope with longer durations of STF, with the former displaying less variability in the number of temporal expression patterns but showing higher amplitudes in terms of individual gene expression. Indeed, intermittent fasting and dietary restriction have been shown to induce different metabolic trade-offs and organ-specific changes in bioenergetics and redox state in mice [36,37].

### Enrichment network reveals key biological processes conserved among the *brain-liver-fats* organ network in the fasted mice

By functionally re-clustering the DEGs, we have also improved the extent of inter-organ overlaps in STF-induced physiological perturbations (Fig. 3a). We identified four highly overlapping organs (LIV, BRN, iBAT, and psWAT) based on their enriched biological pathways (Additional file 7: Table S6). Hereafter, we refer to this organ group as the brain-liver-fats organ network. To further our understanding of the biological implications of STF among this organ network, we performed network enrichment analysis against the Reactome database with the 349 genes integrating the 37 shared GO terms. The network enrichment of 63 reaction pathways (nodes), with 70 connections (edges), resulting from 188 of the 349 genes that passed the significance selection criteria (see Methods), shows that immune-related pathways account for 48% of the total enriched categories, followed by muscle contraction (12.94%) and neuronal system (9.41%) (Fig. 4a,b; Additional file 8: Table S7).

**Figure 4.**
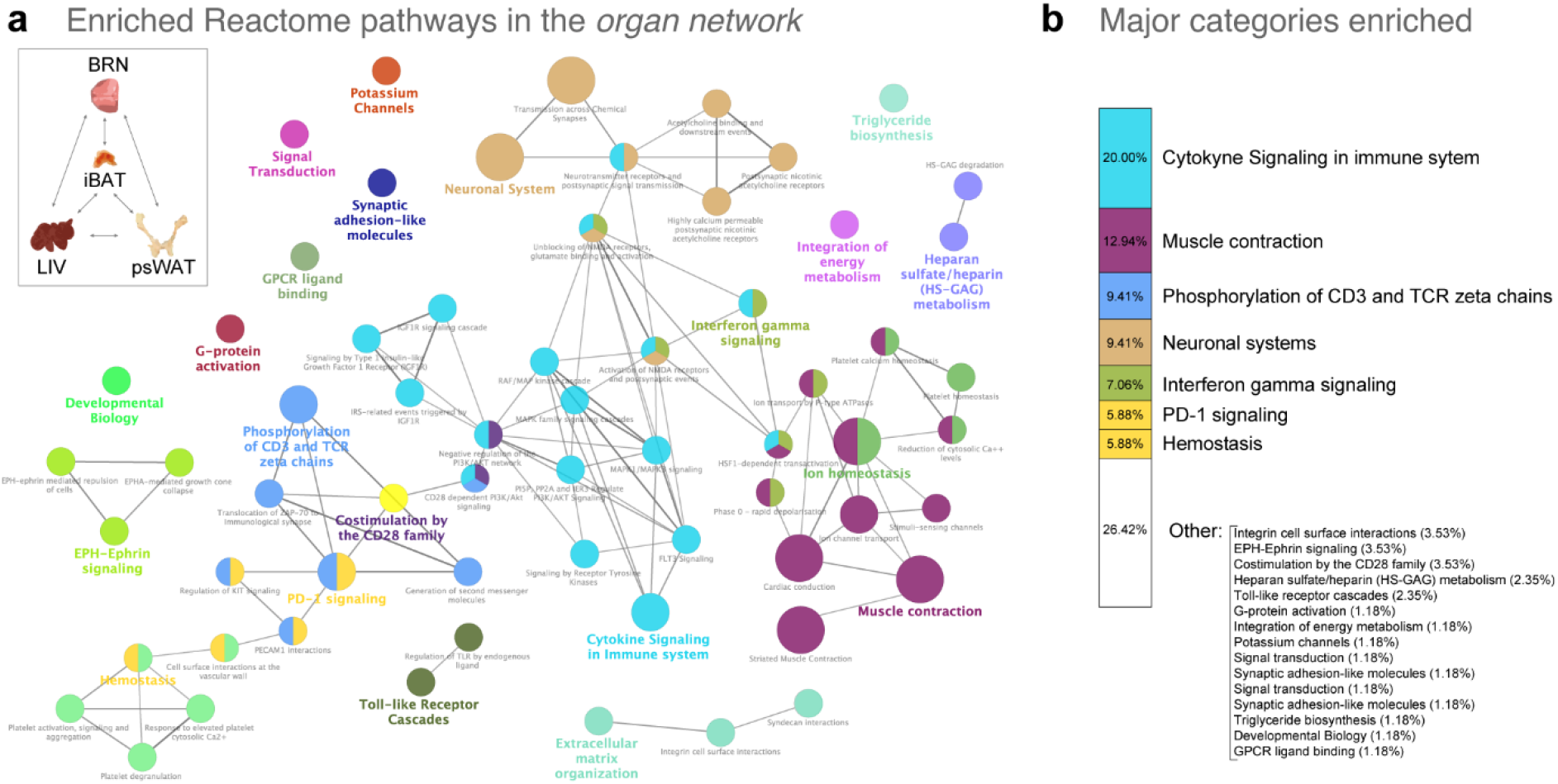
Enrichment network reveals key biological processes conserved among the *brain-liver-fat* network in the fasted mice. **a)** Reactome enrichment network of 63 reaction pathways (nodes) and 70 connections (edges), resulting from 188 of 349 DEGs that were extracted from the 37 overlapping GO terms among BRN, LIV, iBAT and psWAT and passed the significance selection criteria (FDR<0.05). The node color indicates biologically similar reactions and the size reflects the number of genes contributing to the pathway. If the reaction pathway shares 50% or more of the contributing genes, then they are connected by an edge. The representative nodes (based on FDR) are indicated by the colored texts. On the top left, a schematic representation of the four organs composing the *brain-liver-fats* organ network, and all of their possible interactions (bidirectional arrows). **b)** The representative categories and percentages of the Reactome pathways enriched from the 188 genes obtained from the *brain-liver-fats* organ network. The categories and colour are the same as in panel a. Organ acronyms are as in Fig. 1.

To better understand the individual organ responses in the collective network, we represented the gene numbers and proportion of the up-regulated genes associated with each of the summarized Reactome terms across the four organs (Additional file 1: Fig. S3a). Additionally, we performed hierarchical clustering analysis on the gene expression matrix of these 188 genes, to highlight the unique and shared STF-induced expression patterns among the four organs (Additional file 1: Fig. S3b). Although we found the transcriptional programs to be highly specific to organ type, the temporal effect of STF is consistent within and between organs, as demonstrated by the sequential clustering of their respective STF Phases (i.e., Phase I and II cluster closer than to Phase III). Importantly, the differential regulation of the organ DEGs (i.e., percent showing increased expression) indicates that the molecular mechanisms leading to perturbations of the shared biological processes are not the same.

### Biological implications of the conserved DEGs among the *brain-liver-fats* organ network in the fasting mice are in agreement with the current literature

To assess the known biological effects of STF-induced changes among the brain-liver-fats organ network, we used Literature Lab (LitLab™) to perform an association analysis of the 188 DEGs mentioned above with the current body of scientific literature. Briefly, LitLab™ queried the PubMed database (30 million abstracts, January 20 of 2020) for articles associated with the gene list of interest and returned the tagged Medical Subject Headings (MeSHs) with statistical significance. This analysis yielded four MeSH terms clusters that mirrored the four major network-derived pathway categories identified in our previous analysis (Fig. 5a). We then summarized the significant Physiology and Pathway-specific MeSH terms returned and identified associations of the brain-liver-fats organ network with the immune and nervous systems, pain tolerance, and also other pathways and physiological parameters (Additional file 1: Fig. S4a). In sum, the corroborated literature findings support the robustness of our network results, indicating relevant conserved changes in pathway connectivity among the brain-liver-fats web in response to STF.

**Figure 5.**
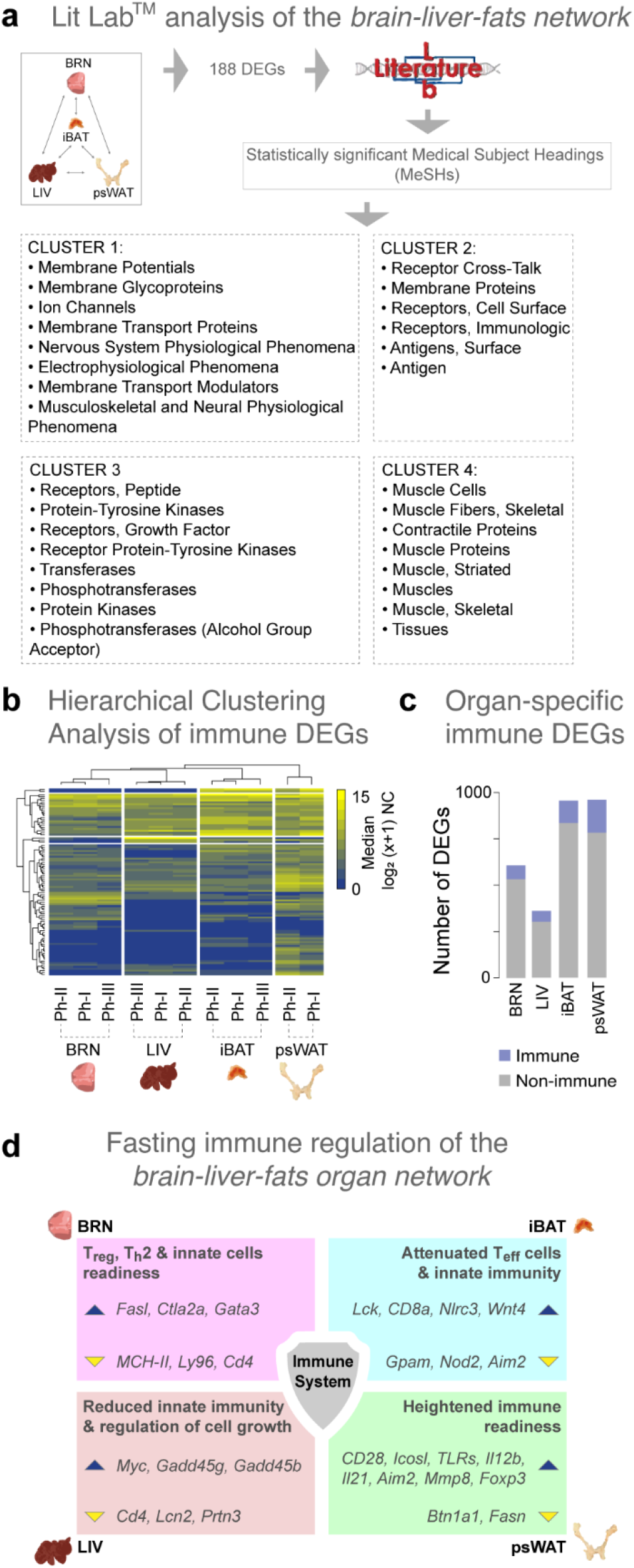
Current understanding of the biological implications of the *brain-liver-fats* organ network in the fasted mice points to immune related processes. **a)** Summary of the significant (cut-off at p-value<0.0228) Medical Subject Heading (MeSH) term clusters associated with the 188 genes among the *brain-liver-fats* organ network were obtained using Literature Lab (LitLabTM). **b)** Gene expression hierarchical clustering analysis of the 96 significant genes (FDR<0.05) from an immune-specific GO enrichment analysis (data not shown) of the 349 genes extracted from the 37 overlapping GO terms from BRN, LIV, iBAT and psWAT. Median mRNA expression levels are represented on a log_2_ (x+1) scale of normalized counts (NC) (0 - not expressed; 15 - highly expressed) against the fasting Phases (Ph). **c)** Proportion of immune-specific genes (blue) queried using the innateDB and Mouse Genome Database, among all DEGs of BRN, LIV, iBAT and psWAT. **d)** The immune system in the context of short-term fasting - a summary schematic highlighting the key changes in the immune regulation of the *brain-liver-fats* organ network during fasting. A brief statement, and example of genes, describing the type of immune change are shown for each organ. The blue and yellow triangles indicate increased and decreased gene expression, respectively. Organ acronyms are as in Fig. 1.

### STF modulates immune-specific transcriptional programs in the brain-liver-fats organ network

We found that among the 349 DEGs extracted from the overlapping GO terms of the brain-liver-fats organ network, 42% (119) are annotated as immune-specific (see Methods). To investigate what aspect of the immune system is centrally involved in the organ network, we re-constructed the enrichment network using an immune-specific GO database (ClueGO). After threshold corrections, 96 genes were retained (Fig. 5b), which resulted in an enrichment of 61 immune-related pathways, with 156 connections (data not shown). The general categories and the proportions of the immune enrichment showed that T-cell regulation processes account for more than 52% of overall processes, followed by leukocyte differentiation (12.5%) and microglial cell activation (9.75%) (Additional file 1: Fig. S4b). The gene numbers and proportion of the up-regulated genes associated with each of the summarized immune-specific GO terms across the four organs (Additional file 1: Fig. S4c).

These results provide a strong indication that T-cell regulation, among other immune processes, are important mediators of STF in the brain-liver-fats organ network. The differential gene expression changes again highlight that the normal immune processes are modulated differently among the four organs composing this organ network.

To better understand the immune-regulated changes in the individual organs of the brain-liver-fats organ network, we asked what are the proportion of immune-related genes among the DEGs for each organ (i.e., the filtered DEGs from the all pair-wise Phase comparisons; Additional file 4: Table S3). Using innateDB and Mouse Genome Database as references, we found that the percentages of immune-specific genes among the DEGs ranged from 11.8% in BRN to 18.8% in psWAT (Fig. 5c). We then identified the top five up and down-regulated immune-related genes among the DEGs from the four organs and evaluated their literature-supported functions (Additional file 1: Fig. S5, Fig. S6). Interestingly, we found that nearly a third (11) of the top 40 differentially expressed genes in the brain-liver-fats organ network have immune-related functions.

Next, we examined the immune-specific genes among the DEGs to infer the immunological status of the four organs. In the psWAT, higher expression of *CD28, Icosl, TLRs, Il12b, Il21, Aim2, Mmp8* (1.5-5.4 log2FC) and lower expression of *Btn1a1* (−4.3 Log2FC) in psWAT suggest a pro-inflammatory activation/environment albeit the absence of infection. Concomitantly, *Foxp3* (1.6 log2FC), which promotes T cell differentiation to Treg, was expressed at higher levels, possibly to prevent overt immune activation. *Fasn* is essential for TLR4 activation in macrophages [38] and its lower expression has been linked to *Nlrp3* inflammasome inhibition and decreased production of the pro-inflammatory cytokine precursor pro-Il1b [39]. Together with *Foxp3* expression, lower expression of *Fasn* (−2.8 log2FC), suggests a regulated heightened state of immune readiness in psWAT during short-term energy loss. We also observed an increased expression of *Cxcr2* (2.36 log2FC), which may be related to reducing adipogenesis [40,41], as a part of the emerging non-conventional functions of chemokine receptors.

In iBAT, *Lck* (promotes CD8 memory T cells) and CD8a (T cell coreceptor for recognition of antigen) showed 2.8 and 3.7 log2FC increases in expression of, respectively. In contrast, *Gpam*, essential for Cd4 T cell metabolic activation [42], was expressed at 2.6 log2FC lower levels. The components of the Nod-like receptor pathway, *Nod2*, and *Aim2* were also expressed at lower levels (−1.6 and −4.4 log2FC, respectively), while *Nlrc3*, an NLR decoy/attenuator shown to attenuate inflammation in myeloid cells, showed increase expression (2.3 log2FC). *Wnt4* has been shown to suppress dendritic cell responsiveness [43] and was increased by 3.3 log2FC. Overall, in iBAT, the effector T cell and innate signaling were reduced, suggesting an anti-inflammatory state during STF.

In BRN, we observed higher expression of FAS ligand (1.9 log2FC) and lower expression (−4.1 log2FC) of *Ly96* (assists immune response via TLR4), suggesting immune-response priming in the absence of MCHII-antigen peptide presentation. Given the increase in *Gata3* (1.9 log2FC), *Cd4* activation was deemed reduced, while T2 differentiation was favored. *Ctla2a* expression also increased (3.2 log2FC), supporting the bias towards memory and regulated immune response in the brain. Collectively, these immune gene changes suggest that in BRN, STF induced the maintenance of higher innate cell activity, likely to preserve organ integrity, via the promotion of less destructive effector mechanisms.

In the liver, the expression of *Cd4* and *Lcn2* (mediators of the innate immune response to bacteria) decreased with fasting time (−1.2 and −3.7 log2FC, respectively). In contrast, *Myc*, which affects cell growth, B cell proliferation, and stem cell renewal, was induced by STF (2.7 log2FC). The Gadd45 family proteins are upregulated under cellular stress [44] and are involved in the activation of S and G2/M checkpoints [45]. Both the gamma and beta forms are critical in the development of pathogenic effector T cells; their deficiency in mice leads to lymphoproliferative syndrome and systemic lupus erythematosus [46]. Downregulation of *Gadd45g* contributes to the pathogenesis of hepatocellular carcinoma in both mouse and human [47]. In T cells, *Gadd45b* is a critical mediator for Th1 response to infection [48]. We observed increases of 2.07 and 1.54 log2FC for *Gadd45g* and *Gadd45b*, respectively, which is likely an attempt to diminish cellular metabolic activities in response to the stress induced by STF. Fasting-induced increases in *Gadd45b* expression is a liver-specific molecular event promoting adaptive metabolic function in mice [49], and are well in line with the role that *Gadd45g* plays as a cold-induced activator of BAT thermogenesis [50].

Finally, to get a second line of evidence that STF affects the transcriptional dynamics of the immune system, we assessed the available literature on the collective topics of fasting-related processes, mouse, gene expression, and immune-related processes (see Methods). This search yielded a total of 241 peer-reviewed articles published over the past ten years. Upon manual inspection, we selected 52 relevant articles for which we then extracted the tagged/associated genes using Gene Retriever™ (Additional file 9: Table S10). Interestingly, the 151 referenced genes that were tagged in more than one article include 10% of the genes in the brain-liver-fats organ network and 23% of the DEGs for BRN, LIV, iBAT, and psWAT. In other words, a small fraction of the immune transcriptional dynamics we observe during STF in mouse has been reported in range of other fasting and diet intervention studies, suggesting that some of the molecular processes triggered by fasting might be independent of its duration. Importantly, this analysis also shows that the vast majority of our findings are novel, thus highlighting the tremendous discovery potential of the experimental approach we used here. In sum, our study expands our current knowledge of molecular processes and pathways shared and modulated in a multi-organ network during short-term energy loss.

## DISCUSSION

Here, we performed mRNA-seq on multiple organs of mice subjected to various periods of STF to understand the molecular mechanisms and biological processes related to short-term energy loss. Our results provide a comprehensive resource on the global mRNA expression changes during STF. We recapitulated some reported physiological/molecular effects from a wide range of longer-term or intermittent fasting studies while providing additional insights into new molecular modulators of fasting in a multi-organ network. Additionally, we present an intuitive analytical method to extrapolate meaningful biological implications from dynamic transcriptomic changes at the organ and systemic levels.

### The analytical approach

Gene expression is highly dynamic, in part due to organ-specific expression patterns and biological variation among individuals [51]. The conventional approach to analyzing dynamic transcriptional responses is to infer the enrichment of biological pathways from gene expression changes at multiple time points by analyzing each time point individually. The implicit assumption that the biological processes are independent of each other - the lack of biological dependency across the time points - limits the ability to pinpoint changes at a pathway level in a biological system [52]. To circumvent this limitation, we utilized HVGs across our dataset as determinants to group samples that best represents their temporal characteristics induced by STF. This data-driven approach allowed us to maximize the detection of the differential temporal gene expression changes that are informative of, but not constrained by, the experimental time points. The latter is of particular importance at the multi-organ level. Given the differential prioritization in energetic re-allocation between organs during short-term energy loss [36,37], it is logical to assume a certain level of asynchrony in their temporal gene expression. Using this approach, we were able to characterize the differential temporal effects of STF on the global gene expression patterns among the nine murine tissues.

Although we observed fold change direction and magnitude in some genes that have been reported in the literature, we failed to observe some known changes, such as a significant increase of *Fgf21* transcript in LIV with increased fasting duration. FGF21 has been shown to act as a negative feedback signal to terminate growth hormone (GH)-stimulated regulation of glucose and lipid metabolism under fasting conditions [53]. In mice, high levels of FGF21 suppresses the activity of GH and reduces the production of insulin-like growth factors (IGF) [54]. In line with the literature, we observed drastic reductions in *Gh* (−4.68 log2FC) and *Igf2* (−3.06 log2FC) expressions in BRN and a trend of increasing raw expression of *Fgf21* in LIV with STF duration (Additional file 2: Table S1). Recently, it was shown that FGF21 is partially required for appropriate gene expression during the fed to fasted transition in mice [55]. FGF21-KO mice and pharmacological blockage of the FGF21 axis did not profoundly disrupt the physiological response to fasting. Also, STF (< 60 hours) did not affect plasma FGF21 level in lean human subjects; however, the mRNA expression of FGF21 receptors (KLB) was decreased in the subcutaneous WAT from both lean and obese subjects [56]. In concordance, we observed a −3.30 log2FC decrease in *Klb* expression in psWAT by STF. Thus, both study heterogeneity and biology may have contributed to these observed gene expression differences in the context of fasting.

An analysis of multiple caloric restriction studies in mice detected relatively few genes that exhibited a consistent expression response across numerous experimental conditions [10]. Thus, relying on specific subsets of DEGs is unlikely to find common biological processes and to provide a meaningful representation of systemic effects. In contrast, a high-level approach, like GO enrichment analysis, are more likely to reveal common biological pathways [57]. Gene set analyses are now the standard practice for functional annotation of gene lists. However, the enrichment bias for multifunctional genes (i.e., frequently represented in GO terms) is an inherent challenge [58], and it drives the generation of biologically non-specific and highly fragile significances in genomic studies [59]. Also, the amount of redundancy and overlaps in GO terms can make result summarization challenging. To address these issues, we clustered the experimentally derived gene list from each organ using the semantic similarity of their functional annotations (i.e., subcDEGs). We then reduced the redundancy of the resulting GO terms using semantic uniqueness, keeping only the most relevant and profoundly affected biological pathways (see Methods). Despite the stringency of our enrichment methods, we obtained increased and sufficient organ overlaps that allowed for the investigation of the biologically relevant events occurring at the multi-organ level.

### A *brain-liver-fats* organ network modulated by STF

We hypothesized that the systemic effects of STF would be, at some level, exerted through biological perturbations shared among multiple organs. We identified four organs that highly overlapped in their enriched biological processes, which we called the brain-liver-fats organ network. Evidence for crosstalk between different pairs of these four organs has been shown in the context of fasting. For example, during prolonged fasting, PPARα and FGF21 signaling between the brain and liver mediates glucose homeostasis [60]. In mice, fasting-induced glycogen shortage activates a liver-brain-adipose neurocircuitry to facilitate fat utilization [61], and the regulation of food intake and glucose homeostasis by liver glycogen is dependent on the hepatic branch of the vagus nerve [62]. In addition, leptin mediated interactions between the brain and adipose depots related to the maintenance of systemic energy balance were recently reviewed [63]. These previous studies provide additional lines of evidence for the existence of our proposed brain-liver-fats organ network in the context of STF.

Within the organ network, we identified immune-related pathways, muscle contraction, neuronal systems, and signal transductions as the top conserved pathways affected by STF. Additionally, with LitLab™, we found strong associations between the genes comprised in our organ network and pain signaling and physiological response to pain. In line with this, fasting and calorie restriction have an analgesic effect in murine models [64–66], and intermittent fasting was proposed as a non-invasive, inexpensive, and implementable strategy to chronic pain treatment (reviewed in [67]). Key underlying mechanisms in fasting-enhanced neuroplasticity include activation of short-term corticosterone increase, reduction in GABAergic inhibition, and increase in protein chaperons and neurotrophic growth factors such as brain-derived neurotrophic factor (BDNF), which exerts positive effects on neuronal survival and synaptogenesis [68–70]. Interestingly, the BDNF pathway was showed a strong association with our organ network gene list, driven by the presence of both *Bdnf* and its receptor, *Ntrk2b* (Additional file 7: Table S6).

As our gene-MeSHs association query goes beyond single-study comparisons (over 30 million abstracts), the results provided unbiased and statistically significant support for our experimental observations. Importantly, we recapitulated several known genes reported in literature associated with fasting, gene expression, immunity, and mouse. As a result of the synthesis of the multi-organ transcriptome, we noted that 90% of the genes encompassing the organ network might represent potential novel molecular modulators of the dynamic biological and immunological perturbations in mice subjected to STF.

### Immune system during homeostatic perturbations

The physiological response to STF is a consortium of organ adaptation, aimed to preserve the most critical functions amidst a systemic decrease in energy availability. The topic of the immune system acting as a regulator of organismal homeostasis in the absence of infection has been recently reviewed [71–73]. Non-infectious signals, such as physiological perturbation (e.g., cold exposure) and diet metabolites, can regulate the equilibrium between different types of immune responses (e.g., intracellular, parasitic, and extracellular) [74,75]. These non-canonical modulations of cytokines in the innate and adaptive immune systems have an essential role in regulating complex organ physiology (reviewed in [72]). These studies suggest that immune cells are well-positioned and equipped to sense homeostatic perturbations and relay signals at the systemic level in the absence of an infection.

In this context, many recent studies have focused on the neuronal regulation of inflammation, neuroimmune circuits in inter-organ communication, and the role immune cells play in the systemic regulation of metabolism and obesity (recently reviewed in [76,77]). For example, macrophage polarization towards a classically (M1-like) activated state is a characteristic of obese adipose tissue [78], and adipose tissue macrophages remain the primary immune participant studied in the context of obesity since their discovery [79,80]. Non-canonical pathways of macrophage activation via metabolites (i.e., glucose, insulin, and palmitate) also result in a continuum of proinflammatory phenotypes [77]. Moreover, sympathetic neuron-associated macrophages were recently identified and shown to affect norepinephrine (NE)-mediate regulation of thermogenesis of adipose tissue by facilitating NE clearance and shifting to a more pro-inflammatory state [81]. Furthermore, liver macrophages contribute to insulin resistance independently of their inflammatory status, via the secretion of IGFBP7, a non-inflammatory factor with a high capacity to bind the insulin receptor and induce lipogenesis and gluconeogenesis through the activation of ERK signaling [82].

### The immune system in the context of short-term fasting

The brain-liver-fats organ network described in this study highlights the importance of the immune processes modulated by STF. However, the mechanisms underlying fasting-induced effects on the immune system remain largely unknown and only recently started to be elucidated [83]. Three recent studies demonstrated that monocytes, naïve B cells, and memory CD8 T cells use bone marrow as a refuge during periods of energy reduction to maintain systemic immune-responsiveness [4–6]. These studies also provided new insights into the integrated immunometabolic response in a state of energy deprivation.

Among the organ network, we found increased expression of genes that negatively regulate monocyte and macrophage activation (*Tiff2* and *Myc*), promote regulatory T cell differentiation (*Ctla2a*), B cell differentiation in the bone marrow (*Fzd9*), cytotoxic T cell differentiation (*Cd8a*), suppression of type 2 immunity and inflammation (Wnt4), and modulation of neuroinflammation and priming *of* innate immunity (*S100a8* and *S100a9*). Moreover, we found that the immune genes showing lower expression values with higher durations of fasting time, are involved in the promotion of inflammation (*Ly96*), activation of the innate immune response (*lfi203* and *Lcn2*), negative regulation of T cell proliferation (*Btn1a1*), inhibition of innate immune response to virus infection (*Trim29*), and mediation of inflammasome activation (*Aim2*). Intuitively, the direction or amplitude of the immunological responses to STF in different organs is unlikely to be the same. Interestingly, we found enriched expression for genes contributing to T cell and the innate response in psWAT, but lower expression levels for inflammation-related genes in iBAT.

Overall, we observed significant increases in the expression of genes involved in T and B cell differentiation and proliferation, suggesting these immune cells are differentiated within the organ network or in circulation. Decreased expression levels of inflammatory markers support a systemic effort to reduce innate immune signals to the adaptive, possibly by blocking cytokine signals and antigen presentation. Intermittent fasting has been shown to alter T cells differentiation bias in the gut, reducing IL-17 producing T cells and increasing regulatory T cells [84]. Fasting-mimicking diets lessen the severity and symptoms in a multiple sclerosis mouse model and are associated with increased regulatory T cell and reduced levels of pro-inflammatory cytokines, Th1 and Th17 cells, and antigen-presenting cells [85]. In line with the literature, the observed changes in our multi-organ immune-transcriptome are such that the effect of STF regroups and reconsolidates the various immune cells, allowing a spectrum of cellular differentiation to occur but restricting immediate reactivity. In sum, our study provides evidence of a consortium of organ adaptation to short-term energy deprivation, in which the immune system plays a central role. Furthermore, we added insights to the molecular events of fasting-induced priming of T cell-mediated immunity, underlining a putative multi-organ effort to support, at the transcriptome level, the recently reported egress of T and B cells to the bone marrow during periods of systemic energy reduction [5,6].

## CONCLUSIONS

While our study was not purposed to decipher the molecular communication between organs or to investigate the migratory behaviors or composition of immune cells under fasting conditions [4–6], our study highlights the centrality of immune-transcriptomic modulations during STF. Using a combinatorial data analysis approach, we provide evidence for the existence of an organ network, formed by the similarities of their biological processes, and the prominent role of the immune system in sensing and modulating systemic homeostatic perturbations in the absence of infection. Additionally, we provide a valuable transcriptome resource to further expand upon our current knowledge of the molecular events occurring across multiple organs during short term energy loss.

## METHODS

### Mice and short-term fasting experiments

All animals used were adult (8-9 weeks of age) male C57BL/6J group-housed mice, on a 12:12h light: dark schedule (lights on at 07:30). For the short-term fasting (STF) experiment, mice were fasted for 2, 8, 12, 18, or 22 hours (n=3 per time point), while the control group (i.e., 0 hour) was fed ad libitum (Fig. 1a). All mice had access to water. All animals were sacrificed by cervical dislocation after the start of the dark cycle (between 21:00 and 21:30), and the following nine organs were collected and processed for mRNA-seq: olfactory bulb (OB), brain (BRN, which includes the telencephalon and diencephalon), cerebellum (CBL), brainstem (BST, which consists of the mesencephalon, pons, and myelencephalon), stomach (STM), liver (LIV), interscapular brown adipose tissue (iBAT), perigonadal white adipose tissue (pgWAT), and posterior-subcutaneous white adipose tissue (psWAT). The organs were immediately frozen and kept at −80°C until further processing.

### RNA extraction and RNA-sequencing

Organs (OB, BRN, CBL, BST, STM, LIV, and iBAT) were homogenized in Lysis RLT Buffer (Qiagen) supplemented with 1% β-mercaptoethanol (Sigma-Aldrich) using the OMNI tissue homogenizer (TH) (OMNI International). Homogenized organ lysates were then loaded onto the QIAshredder homogenizer spin columns (Qiagen) for further homogenization and elimination of insoluble debris. Total RNA was extracted using RNeasy Mini Kit (Qiagen), according to the manufacturer’s protocol. White-adipose tissues (pgWAT and psWAT) were homogenized in Qiazol (Qiagen) and RNA extracted with the Lipid RNeasy Lipid Tissue Mini Kit (Qiagen). mRNA was prepared for sequencing using the TruSeq stranded mRNA sample preparation kit (Illumina), with a selected insert size of 120–210 bp. All samples were sequenced on an Illumina HiSeq 4000, to generate paired-end 150 bp sequencing reads and at an average depth of 39.98 ± 1.05 (SEM) million reads per sample (Additional file 2: Table S1).

### Short reads alignment to reference genome and transcriptome

The quality of the reads was assessed using FastQC (KBase). Sample Fastq files were aligned to the mouse reference genome mm10/GRCm38.p5 using TopHat2 [86] with 2 mismatches allowed. Reads were retained only if uniquely mapped to the genome. We used HTSeq-count (0.9.1, −t exon and –m union) to obtain the number of reads that were mapped to each gene in Gencode M16. Bigwig files were generated from bam files for visualization using RSeQC [87].

### Pre-Processing

The schematic of the bioinformatic workflow is presented in Additional file 1: Fig. S1. Prior to the analysis, the mapped read counts were filtered for annotated genes using org.Mm.eg.db [88]. A count per million threshold equivalent to ~10 raw expression value was applied to remove all lowly expressed genes, and only genes having ≥ 3 samples above the threshold were kept. Samples with total reads lower than two standard deviations from the tissue means were removed.

### Clustering group calls and bootstrap validations

Highly variable genes (HVGs) were queried from normalized log2 expression values (log2 (x+1) NC) of the filtered datasets for informative genes and were used for the data-driven clustering calls. Briefly, we calculated the gene-specific variance and regressed against its mean log-expression value and applied a false discovery rate (FDR) <0.05 was applied to denote significance and resulted in between 829-1190 HVGs per organ. Spearman’s Rho was then used to calculate the correlation distance matrix among these genes per organ:

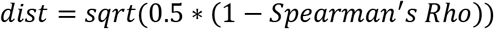

To determine the optimal number of phase clusters, hierarchical clustering (method = complete) was performed on the correlation distance matrixes of the HVGs and bootstrapped to 5000 iterations. A bootstrap mean cutoff (≥0.65) was used to determine the significance of the cluster fit, and any clusters not meeting the criteria were grouped with the previous node. A cluster size of three was found to be optimal for most organs, which we designated as Phases I, II, and III (Additional file 3: Table S2).

### Determining the differentially expressed genes

Differential gene expression analysis was performed independently for each organ between all Phase groups using the following criteria: log2FC >1.0 and FDR <0.05 in DESeq2 (Additional file 4: Table S3). To reduce complexity, only the most extreme (top and bottom 12.5%) of the log2FC ranked DEGs were kept for organs with >500 unique DEGs for all subsequent analyses.

### Gene ontology enrichment analysis

Several strategies were used to perform the gene functional enrichment analysis to maximize annotation coverage and to provide meaningful interpretations for the DEGs. Briefly, the unique DEGs from each organ were submitted, as a single gene set, to query for Mus musculus GO terms associated with biological processes using the following algorithms and databases: gene over-representation test (enrichGO) and gene set enrichment analysis, both of which are provided by ClusterProfiler [89], and the Database for Annotation, Visualization and Integrated Discovery (DAVID, [90]). The minimal gene set size was set at three, and an FDR<0.05 cutoff was set for significance. When applicable, the number of permutations was set at 1000. To manage the output size, a cutoff of 150 GO terms, ranked by FDR, was implemented. All GO enrichment results were semantically reduced using REVIGO, with a dispensability threshold set at <0.4 [91].

### Functional re-clustering of DEGs - subcDEGs

From the unique DEGs obtained from all pairwise comparisons, semantic similarity measures were calculated to determine the gene function-based clusters in each organ [92] (Additional file 7: Table S6). The optimal numbers of clusters were determined using dynamicTreeCut with deepSplit = 0 [91]. The gene function-based clusters of DEGs are referred to as the sub-clustered DEGs (subcDEGs). The SubcDEGS were then re-submitted for GO enrichment analysis and reduced for term redundancy as described above. SubcDEGS that did not yield GO enrichment were excluded from downstream analysis.

To evaluate the changes in gene expression pattern through the different STF phases, we calculated the log2FCs (against 0 hour) of the functional gene clusters mentioned above. Two-sided Wilcoxon Rank test was used to determine the statistical significance between means at p≤0.05. The mean variances of the log2FCs were regressed against the fasting phases (Spearman’s correlation was used).

### Gene expression pattern mining

Unique patterns across the STF phases were determined from log2(x+1) NC of the genes that composed all the subcDEGs of each organ and that yielded GO enrichment. Pattern mining was performed using topology overlap matrix-based dissimilarity algorithms (WGCNA), followed by the optimal cluster numbers determined using dynamicTreeCut with deepSplit = 0.

### Network enrichment analysis

A list of 349 unique genes was extracted from the 37 GO terms shared among the top four overlapping organs: BRN, LIV, iBAT, and psWAT, to further investigate the enriched biological processes in the brain-liver-fats organ network observed in the fasted mice (Additional file 7: Table S6). The Reactome database [93] and ClueGO [94] were used to determine and visualize the enrichment networks and the conserved protein pathways. The following parameters were used to construct the enrichment network: min/max GO level = 3-20, Number of Genes = 3, Min Percentage = 3, Kappa Score Threshold = 0.4, Sharing Group Percentage = 50 and the statistical significance was set at FDR<0.05.

To evaluate the immune components of the organs in the brain-liver-fats organ network (BRN, LIV, iBAT, and psWAT), we retrieved a comprehensive list of immune-related genes in mice from innateDB [95] and Mouse Genome Database (http://www.informatics.jax.org/vocab/gene_ontology/GO:0002376) [96]. A compiled list of 3022 genes was used to identify the immune-related genes in this study.

### Literature Mining - Acumenta Literature Lab (LitLabTM)

LitLab™ (Acumenta Biotech) allows the identification of biological and biochemical terms that are significantly associated with the literature from a gene set. The analysis provides additional meaning to experimentally derived gene and protein data [97]. The LitLab™ database contains current gene, biological, and biochemical references in every indexed PubMed abstract and are updated quarterly (currently at 30 million). LitLab™ calculates the frequencies of the input genes with each of 86,000 terms in the Literature Lab™ database (as of January 20th, 2020) and compare the values with that of a 1000 random gene sets to determine statistical significance (p-value<0.0228). LitLab™ is composed of four main applications: Term Viewer, PLUS, Editor, and Gene Retriever.

### PubMed-based literature searches

A literature search of the PubMed databased on “fasting” and “immunity” in mice was conducted using the following search strategy: (“intermittent fasting”[All Fields] OR “Caloric Restriction”[Mesh] OR “Food Deprivation”[All] OR “food restriction”[All Fields] OR “fasting”[MeSH Terms] OR “fasting”[All Fields]) AND (“gene expression”[All Fields] OR “Gene Expression”[Mesh] OR “transcriptome”[All Fields] OR “transcriptome”[MeSH Terms]) AND (“mice”[MeSH Terms] OR “mice”[All Fields]) AND (“2010/01/20”[PDAT] : “2020/01/20”[PDAT]) AND (“Immune System Phenomena”[Mesh] OR “Immunity”[Mesh] OR “Immune System”[Mesh] OR immune[All Fields] OR “immune response”[All Fields] OR “inflammation”[MeSH Terms] OR “inflammation”[All Fields] OR “infection”[All Fields]).

To further evaluate our findings with the current literature, we extracted the gene referenced from the relevant resultant articles using Gene Retriever™ (Acumenta). Gene Retriever™ is a data mining solution to retrieve all genes associated with a list of PubMed articles. Gene Retriever processes an input list of PubMed IDs and produces an analysis of the genes mentioned in the title, text, and MeSH tags of each record. Results are then statistically ranked and presented in a spreadsheet to enable quick and comprehensive analyses. Hyperlinks are added within the spreadsheet to enable instant review of the genes or PubMed IDs of interest (Additional file 9: Table S8).

### Software

All analyses and graphics were performed and generated in R unless otherwise stated. ClueGO was performed in Cytoscape [98]. Adobe Illustrator (Adobe Inc.) was used to prepare the final figures.

## ABBREVIATIONS

STF: Short-term fasting
OB: olfactory bulb
BRN: brain
CBL: cerebellum
BST: brainstem
STM: stomach
LIV: liver
iBAT: interscapular brown adipose tissue
pgWAT: perigonadal white adipose tissue
psWAT: posterior-subcutaneous white adipose tissue
HVG: highly-variable genes
log2(x+1) NC: log2 expression values
HCA: hierarchical clustering analysis
FDR: false discovery rate
DEGs: differentially expressed genes
FC: fold change
log2FC: log2 fold change
GO: Gene Ontology
subcDEGs: sub-clustered DEGs
LitLabTM: Acumenta Literature Lab
MeSH: Medical Subject Headings
GH: growth hormone
IGF: insulin-like growth factors
BDNF: brain-derived neurotropic factor
NE: norepinephrine

## DECLARATIONS

### Ethics approval and consent to participate

The use and care of animals used in this study were in accordance with UK Home Office regulations and under a project license approved by the Wellcome Sanger Institute Animal Welfare and Ethical Review Body.

### Consent for publication

The authors have consented the study for publication.

### Availability of data and materials

The raw RNA-seq data from the 162 samples will be available at GEO with the peer-reviewed publication of this study.

## AUTHOR’S CONTRIBUTIONS

LRS, MG, and SSY designed and led the study. LRS, EHA, LSM, KW, MCL performed experiments. SSYH, MM, MLRTS, AS, MG, and LRS analyzed data and SSYH, MG, and LRS interpreted data. DWL, and DC contributed analysis tools and interpreted data. SSYH, MG, and LRS wrote and MM, AS, DC and DWL commented on the manuscript. All authors have read and approved the final manuscript.

## ACKNOWLEDGMENTS

We would like to acknowledge the Clinical Genomic Core of Sidra Medicine for performing the mRNA-seq, Diogo Manoel and the other Saraiva Lab members for their insightful inputs on data analysis and comments on the manuscript.

## SUPPLEMENTARY INFORMATION

**Additional file 1: Figure S1-S6.**

**Additional file 2: Table S1.** Gene expression estimates as normalized counts (NCs) for all filtered genes and all samples.

**Additional file 3: Table S2.** Phase cluster analysis results based on the high variable genes (HVGs).

**Additional file 4: Table S3.** Differential expression analysis for all pairwise Phase comparisons for OB, BRN, CBL, BST, STM, LIV, iBAT, and psWAT.

**Additional file 5: Table S4.** Gene expression estimates as a log_2_ (x+1) normalized counts (NCs) for all DEGs for all samples. Median, mean and SEM are also displayed.

**Additional file 6: Table S5.** Gene ontology (GO) enrichment for all subcDEGs in all organs.

**Additional file 7: Table S6.** The overlapping enriched gene ontology (GO) terms for all subcDEGs that were used for the *brain-liver-fats* organ network analysis.

**Additional file 8: Table S7.** Enriched categories for the Reactome pathway enrichment of the *brain-liver-fats* organ network.

**Additional file 9: Table S8.** Gene list compiled from 52 triaged PubMed articles, using Gene Retriever™.

## REFERENCES

1. Longo VD, Panda S. Fasting, Circadian Rhythms, and Time-Restricted Feeding in Healthy Lifespan. Cell Metab. 2016;23:1048–59.

2. Mattson MP, Longo VD, Harvie M. Impact of intermittent fasting on health and disease processes. Ageing Res Rev. 2017;39:46–58.

3. Stockman M-C, Thomas D, Burke J, Apovian CM. Intermittent Fasting: Is the Wait Worth the Weight? Curr Obes Rep. 2018;7:172–85.

4. Nagai M, Noguchi R, Takahashi D, Morikawa T, Koshida K, Komiyama S, et al. Fasting-Refeeding Impacts Immune Cell Dynamics and Mucosal Immune Responses. Cell. 2019;178:1072–1087.e14.

5. Collins N, Han S-J, Enamorado M, Link VM, Huang B, Moseman EA, et al. The Bone Marrow Protects and Optimizes Immunological Memory during Dietary Restriction. Cell. 2019;178:1088–1101.e15.

6. Jordan S, Tung N, Casanova-Acebes M, Chang C, Cantoni C, Zhang D, et al. Dietary Intake Regulates the Circulating Inflammatory Monocyte Pool. Cell. 2019;178:1102–1114.e17.

7. Intermountain Health Care, Inc. The Fasting II Study [Internet]. 2017 [cited 2020 Mar 5]. Available from: https://clinicaltrials.gov/ct2/show/NCT01792986

8. Kessler CS, Stange R, Schlenkermann M, Jeitler M, Michalsen A, Selle A, et al. A nonrandomized controlled clinical pilot trial on 8 wk of intermittent fasting (24 h/wk). Nutr Burbank Los Angel Cty Calif. 2018;46:143–152.e2.

9. Stekovic S, Hofer SJ, Tripolt N, Aon MA, Royer P, Pein L, et al. Alternate Day Fasting Improves Physiological and Molecular Markers of Aging in Healthy, Non-obese Humans. Cell Metab. 2019;30:462–476.e5.

10. Swindell WR. Genes and gene expression modules associated with caloric restriction and aging in the laboratory mouse. BMC Genomics. 2009;10:585.

11. Kim K-H, Kim YH, Son JE, Lee JH, Kim S, Choe MS, et al. Intermittent fasting promotes adipose thermogenesis and metabolic homeostasis via VEGF-mediated alternative activation of macrophage. Cell Res. 2017;27:1309–26.

12. Goldstein I, Baek S, Presman DM, Paakinaho V, Swinstead EE, Hager GL. Transcription factor assisted loading and enhancer dynamics dictate the hepatic fasting response. Genome Res. 2017;27:427–39.

13. Kinouchi K, Magnan C, Ceglia N, Liu Y, Cervantes M, Pastore N, et al. Fasting Imparts a Switch to Alternative Daily Pathways in Liver and Muscle. Cell Rep. 2018;25:3299–3314.e6.

14. Rennert C, Vlaic S, Marbach-Breitrück E, Thiel C, Sales S, Shevchenko A, et al. The Diurnal Timing of Starvation Differently Impacts Murine Hepatic Gene Expression and Lipid Metabolism – A Systems Biology Analysis Using Self-Organizing Maps. Front Physiol. 2018;9.

15. Derous D, Mitchell SE, Green CL, Wang Y, Han JDJ, Chen L, et al. The Effects of Graded Levels of Calorie Restriction: X. Transcriptomic Responses of Epididymal Adipose Tissue. J Gerontol A Biol Sci Med Sci. 2018;73:279–88.

16. Ng GY-Q, Kang S-W, Kim J, Alli-Shaik A, Baik S-H, Jo D-G, et al. Genome-Wide Transcriptome Analysis Reveals Intermittent Fasting-Induced Metabolic Rewiring in the Liver. Dose-Response. 2019;17:1559325819876780.

17. Smith RL, Soeters MR, Wüst RCI, Houtkooper RH. Metabolic Flexibility as an Adaptation to Energy Resources and Requirements in Health and Disease. Endocr Rev. 2018;39:489–517.

18. Jensen TL, Kiersgaard MK, Sørensen DB, Mikkelsen LF. Fasting of mice: a review. Lab Anim. 2013;47:225–40.

19. Hakvoort TBM, Moerland PD, Frijters R, Sokolović A, Labruyère WT, Vermeulen JLM, et al. Interorgan Coordination of the Murine Adaptive Response to Fasting. J Biol Chem. 2011;286:16332–16343.

20. Schupp M, Chen F, Briggs ER, Rao S, Pelzmann HJ, Pessentheiner AR, et al. Metabolite and transcriptome analysis during fasting suggest a role for the p53-Ddit4 axis in major metabolic tissues. BMC Genomics. 2013;14:758.

21. Pan Q, Shai O, Misquitta C, Zhang W, Saltzman AL, Mohammad N, et al. Revealing Global Regulatory Features of Mammalian Alternative Splicing Using a Quantitative Microarray Platform. Mol Cell. 2004;16:929–41.

22. Su AI, Wiltshire T, Batalov S, Lapp H, Ching KA, Block D, et al. A gene atlas of the mouse and human protein-encoding transcriptomes. Proc Natl Acad Sci. 2004;101:6062–7.

23. Zhang W, Morris QD, Chang R, Shai O, Bakowski MA, Mitsakakis N, et al. The functional landscape of mouse gene expression. J Biol. 2004;3:21.

24. Caba M, Pabello M, Moreno ML, Meza E. Main and accessory olfactory bulbs and their projections in the brain anticipate feeding in food-entrained rats. Chronobiol Int. 2014;31:869–77.

25. Mercer SW, Williamson DH. The influence of starvation and natural refeeding on the rate of triacylglycerol/fatty acid substrate cycling in brown adipose tissue and different white adipose sites of the rat in vivo. The role of insulin and the sympathetic nervous system. Biosci Rep. 1988;8:147–53.

26. Nolasco N, Juárez C, Morgado E, Meza E, Caba M. A Circadian Clock in the Olfactory Bulb Anticipates Feeding during Food Anticipatory Activity. PLOS ONE. 2012;7:e47779.

27. Palou M, Sánchez J, Priego T, Rodríguez AM, Picó C, Palou A. Regional differences in the expression of genes involved in lipid metabolism in adipose tissue in response to short- and medium-term fasting and refeeding. J Nutr Biochem. 2010;21:23–33.

28. Schoettl T, Fischer IP, Ussar S. Heterogeneity of adipose tissue in development and metabolic function. J Exp Biol. 2018;221.

29. Lambrecht NWG, Yakubov I, Sachs G. Fasting-induced changes in ECL cell gene expression. Physiol Genomics. American Physiological Society; 2007;31:183–92.

30. Sokolović M, Sokolović A, Wehkamp D, van Themaat EVL, de Waart DR, Gilhuijs-Pederson LA, et al. The transcriptomic signature of fasting murine liver. BMC Genomics. 2008;9:528.

31. Tang H-N, Tang C-Y, Man X-F, Tan S-W, Guo Y, Tang J, et al. Plasticity of adipose tissue in response to fasting and refeeding in male mice. Nutr Metab. 2017;14:3.

32. Wu V, Sumii K, Tari A, Sumii M, Walsh JH. Regulation of rat antral gastrin and somatostatin gene expression during starvation and after refeeding. Gastroenterology. 1991;101:1552–8.

33. Bolshakova N, Azuaje F, Cunningham P. A knowledge-driven approach to cluster validity assessment. Bioinformatics. 2005;21:2546–7.

34. Wolting C, McGlade CJ, Tritchler D. Cluster analysis of protein array results via similarity of Gene Ontology annotation. BMC Bioinformatics. 2006;7:338.

35. Chi Y, Youn DY, Xiaoli AM, Liu L, Pessin JB, Yang F, et al. Regulation of gene expression during the fasting–feeding cycle of the liver displays mouse strain specificity. J Biol Chem. 2020;jbc.RA119.012349.

36. Książek A, Konarzewski M. Effect of dietary restriction on immune response of laboratory mice divergently selected for basal metabolic rate. Physiol Biochem Zool PBZ. 2012;85:51–61.

37. Chausse B, Vieira-Lara MA, Sanchez AB, Medeiros MHG, Kowaltowski AJ. Intermittent Fasting Results in Tissue-Specific Changes in Bioenergetics and Redox State. PLOS ONE. 2015;10:e0120413.

38. Carroll RG, Zasłona Z, Galván-Peña S, Koppe EL, Sévin DC, Angiari S, et al. An unexpected link between fatty acid synthase and cholesterol synthesis in proinflammatory macrophage activation. J Biol Chem. 2018;293:5509–21.

39. Moon J-S, Lee S, Park M-A, Siempos II, Haslip M, Lee PJ, et al. UCP2-induced fatty acid synthase promotes NLRP3 inflammasome activation during sepsis. J Clin Invest. 2015;125:665–80.

40. Dyer DP, Nebot JB, Kelly CJ, Medina-Ruiz L, Schuette F, Graham GJ. The chemokine receptor CXCR2 contributes to murine adipocyte development. J Leukoc Biol. 2019;105:497–506.

41. Kusuyama J, Komorizono A, Bandow K, Ohnishi T, Matsuguchi T. CXCL3 positively regulates adipogenic differentiation. J Lipid Res. 2016;57:1806–20.

42. Faris R, Fan Y-Y, De Angulo A, Chapkin RS, deGraffenried LA, Jolly CA. Mitochondrial glycerol-3-phosphate acyltransferase-1 is essential for murine CD4(+) T cell metabolic activation. Biochim Biophys Acta. 2014;1842:1475–82.

43. Hung L-Y, Johnson JL, Ji Y, Christian DA, Herbine KR, Pastore CF, et al. Cell-Intrinsic Wnt4 Influences Conventional Dendritic Cell Fate Determination to Suppress Type 2 Immunity. J Immunol Baltim Md 1950. 2019;203:511–9.

44. Takekawa M, Saito H. A family of stress-inducible GADD45-like proteins mediate activation of the stress-responsive MTK1/MEKK4 MAPKKK. Cell. 1998;95:521–30.

45. Vairapandi M, Balliet AG, Hoffman B, Liebermann DA. GADD45b and GADD45g are cdc2/cyclinB1 kinase inhibitors with a role in S and G2/M cell cycle checkpoints induced by genotoxic stress. J Cell Physiol. 2002;192:327–38.

46. Liu L, Tran E, Zhao Y, Huang Y, Flavell R, Lu B. Gadd45 beta and Gadd45 gamma are critical for regulating autoimmunity. J Exp Med. 2005;202:1341–7.

47. Zhang L, Yang Z, Ma A, Qu Y, Xia S, Xu D, et al. Growth arrest and DNA damage 45G down-regulation contributes to Janus kinase/signal transducer and activator of transcription 3 activation and cellular senescence evasion in hepatocellular carcinoma. Hepatol Baltim Md. 2014;59:178–89.

48. Lu B, Ferrandino AF, Flavell RA. Gadd45beta is important for perpetuating cognate and inflammatory signals in T cells. Nat Immunol. 2004;5:38–44.

49. Fuhrmeister J, Zota A, Sijmonsma TP, Seibert O, Cıngır Ş, Schmidt K, et al. Fasting-induced liver GADD45β restrains hepatic fatty acid uptake and improves metabolic health. EMBO Mol Med. 2016;8:654–69.

50. Gantner ML, Hazen BC, Conkright J, Kralli A. GADD45γ regulates the thermogenic capacity of brown adipose tissue. Proc Natl Acad Sci U S A. 2014;111:11870–5.

51. Tan MH, Li Q, Shanmugam R, Piskol R, Kohler J, Young AN, et al. Dynamic landscape and regulation of RNA editing in mammals. Nature. 2017;550:249–54.

52. Khatri P, Sirota M, Butte AJ. Ten Years of Pathway Analysis: Current Approaches and Outstanding Challenges. PLOS Comput Biol. 2012;8:e1002375.

53. Chen W, Hoo RL, Konishi M, Itoh N, Lee P-C, Ye H, et al. Growth hormone induces hepatic production of fibroblast growth factor 21 through a mechanism dependent on lipolysis in adipocytes. J Biol Chem. 2011;286:34559–66.

54. Inagaki T, Lin VY, Goetz R, Mohammadi M, Mangelsdorf DJ, Kliewer SA. Inhibition of Growth Hormone Signaling by the Fasting-Induced Hormone FGF21. Cell Metab. 2008;8:77–83.

55. Antonellis PJ, Hayes MP, Adams AC. Fibroblast Growth Factor 21-Null Mice Do Not Exhibit an Impaired Response to Fasting. Front Endocrinol [Internet]. 2016 [cited 2020 Jan 15];7. Available from: https://www.frontiersin.org/articles/10.3389/fendo.2016.00077/full

56. Nygaard EB, Ørskov C, Almdal T, Vestergaard H, Andersen B. Fasting decreases plasma FGF21 in obese subjects and the expression of FGF21 receptors in adipose tissue in both lean and obese subjects. J Endocrinol. 2018;239:73–80.

57. Subramanian A, Tamayo P, Mootha VK, Mukherjee S, Ebert BL, Gillette MA, et al. Gene set enrichment analysis: a knowledge-based approach for interpreting genome-wide expression profiles. Proc Natl Acad Sci U S A. 2005;102:15545–50.

58. Gillis J, Pavlidis P. The impact of multifunctional genes on “guilt by association” analysis. PloS One. 2011;6:e17258.

59. Ballouz S, Pavlidis P, Gillis J. Using predictive specificity to determine when gene set analysis is biologically meaningful. Nucleic Acids Res. 2017;45:e20–e20.

60. Liang Q, Zhong L, Zhang J, Wang Y, Bornstein SR, Triggle CR, et al. FGF21 Maintains Glucose Homeostasis by Mediating the Cross Talk Between Liver and Brain During Prolonged Fasting. Diabetes. 2014;63:4064–75.

61. Izumida Y, Yahagi N, Takeuchi Y, Nishi M, Shikama A, Takarada A, et al. Glycogen shortage during fasting triggers liver–brain–adipose neurocircuitry to facilitate fat utilization. Nat Commun. 2013;4:1–9.

62. López-Soldado I, Fuentes-Romero R, Duran J, Guinovart JJ. Effects of hepatic glycogen on food intake and glucose homeostasis are mediated by the vagus nerve in mice. Diabetologia. 2017;60:1076–83.

63. Caron A, Lee S, Elmquist JK, Gautron L. Leptin and brain–adipose crosstalks. Nat Rev Neurosci. 2018;19:153–65.

64. Hargraves WA, Hentall ID. Analgesic effects of dietary caloric restriction in adult mice. Pain. 2005;114:455–61.

65. Liu Y, Ni Y, Zhang W, Sun Y-E, Ma Z, Gu X. Antinociceptive effects of caloric restriction on post-incisional pain in nonobese rats. Sci Rep. 2017;7:1805.

66. Lee J-Y, Lee GJ, Lee PR, Won CH, Kim D, Kang Y, et al. The analgesic effect of refeeding on acute and chronic inflammatory pain. Sci Rep. 2019;9.

67. Sibille KT, Bartsch F, Reddy D, Fillingim RB, Keil A. Increasing Neuroplasticity to Bolster Chronic Pain Treatment: A Role for Intermittent Fasting and Glucose Administration? J Pain Off J Am Pain Soc. 2016;17:275–81.

68. Spolidoro M, Baroncelli L, Putignano E, Maya-Vetencourt JF, Viegi A, Maffei L. Food restriction enhances visual cortex plasticity in adulthood. Nat Commun. 2011;2:320.

69. Mattson MP, Duan W, Guo Z. Meal size and frequency affect neuronal plasticity and vulnerability to disease: cellular and molecular mechanisms. J Neurochem. 2003;84:417–31.

70. van Praag H, Fleshner M, Schwartz MW, Mattson MP. Exercise, energy intake, glucose homeostasis, and the brain. J Neurosci Off J Soc Neurosci. 2014;34:15139–49.

71. Caputa G, Castoldi A, Pearce EJ. Metabolic adaptations of tissue-resident immune cells. Nat Immunol. Nature Publishing Group; 2019;20:793–801.

72. Rankin LC, Artis D. Beyond Host Defense: Emerging Functions of the Immune System in Regulating Complex Tissue Physiology. Cell. 2018;173:554–67.

73. Veiga-Fernandes H, Freitas AA. The S(c)ensory Immune System Theory. Trends Immunol. 2017;38:777–88.

74. Shibata N, Kunisawa J, Kiyono H. Dietary and Microbial Metabolites in the Regulation of Host Immunity. Front Microbiol. 2017;8.

75. Macpherson AJ, de Agüero MG, Ganal-Vonarburg SC. How nutrition and the maternal microbiota shape the neonatal immune system. Nat Rev Immunol. Nature Publishing Group; 2017;17:508–17.

76. Huh JR, Veiga-Fernandes H. Neuroimmune circuits in inter-organ communication. Nat Rev Immunol. 2019;

77. Larabee CM, Neely OC, Domingos AI. Obesity: a neuroimmunometabolic perspective. Nat Rev Endocrinol. 2020;16:30–43.

78. Lumeng CN, Bodzin JL, Saltiel AR. Obesity induces a phenotypic switch in adipose tissue macrophage polarization. J Clin Invest. 2007;117:175–84.

79. Weisberg SP, McCann D, Desai M, Rosenbaum M, Leibel RL, Ferrante AW. Obesity is associated with macrophage accumulation in adipose tissue. J Clin Invest. 2003;112:1796–808.

80. Xu H, Barnes GT, Yang Q, Tan G, Yang D, Chou CJ, et al. Chronic inflammation in fat plays a crucial role in the development of obesity-related insulin resistance. J Clin Invest. 2003;112:1821–30.

81. Pirzgalska RM, Seixas E, Seidman JS, Link VM, Sánchez NM, Mahú I, et al. Sympathetic neuron–associated macrophages contribute to obesity by importing and metabolizing norepinephrine. Nat Med. 2017;23:1309–18.

82. Morgantini C, Jager J, Li X, Levi L, Azzimato V, Sulen A, et al. Liver macrophages regulate systemic metabolism through non-inflammatory factors. Nat Metab. 2019;1:445–59.

83. Buono R, Longo VD. When Fasting Gets Tough, the Tough Immune Cells Get Going—or Die. Cell. 2019;178:1038–40.

84. Cignarella F, Cantoni C, Ghezzi L, Salter A, Dorsett Y, Chen L, et al. Intermittent Fasting Confers Protection in CNS Autoimmunity by Altering the Gut Microbiota. Cell Metab. 2018;27:1222–1235.e6.

85. Choi IY, Piccio L, Childress P, Bollman B, Ghosh A, Brandhorst S, et al. A Diet Mimicking Fasting Promotes Regeneration and Reduces Autoimmunity and Multiple Sclerosis Symptoms. Cell Rep. 2016;15:2136–46.

86. Kim D, Pertea G, Trapnell C, Pimentel H, Kelley R, Salzberg SL. TopHat2: accurate alignment of transcriptomes in the presence of insertions, deletions and gene fusions. Genome Biol. 2013;14:R36.

87. Wang L, Wang S, Li W. RSeQC: quality control of RNA-seq experiments. Bioinformatics. 2012;28:2184–5.

88. Carlson M. org.Mm.eg.db: Genome wide annotation for Mouse [Internet]. 2019. Available from: http://bioconductor.org/packages/release/data/annotation/html/org.Mm.eg.db.html

89. Yu G, Wang L-G, Han Y, He Q-Y. clusterProfiler: an R Package for Comparing Biological Themes Among Gene Clusters. OMICS J Integr Biol. 2012;16:284–7.

90. Huang DW, Sherman BT, Lempicki RA. Bioinformatics enrichment tools: paths toward the comprehensive functional analysis of large gene lists. Nucleic Acids Res. 2009;37:1–13.

91. Supek F, Bošnjak M, Škunca N, Šmuc T. REVIGO Summarizes and Visualizes Long Lists of Gene Ontology Terms. PLOS ONE. 2011;6:e21800.

92. Yu G, Li F, Qin Y, Bo X, Wu Y, Wang S. GOSemSim: an R package for measuring semantic similarity among GO terms and gene products. Bioinformatics. 2010;26:976–8.

93. Croft D, Mundo AF, Haw R, Milacic M, Weiser J, Wu G, et al. The Reactome pathway knowledgebase. Nucleic Acids Res. 2014;42:D472–477.

94. Bindea G, Mlecnik B, Hackl H, Charoentong P, Tosolini M, Kirilovsky A, et al. ClueGO: a Cytoscape plug-in to decipher functionally grouped gene ontology and pathway annotation networks. Bioinforma Oxf Engl. 2009;25:1091–3.

95. Breuer K, Foroushani AK, Laird MR, Chen C, Sribnaia A, Lo R, et al. InnateDB: systems biology of innate immunity and beyond—recent updates and continuing curation. Nucleic Acids Res. 2013;41:D1228–33.

96. Bult CJ, Blake JA, Smith CL, Kadin JA, Richardson JE, Mouse Genome Database Group. Mouse Genome Database (MGD) 2019. Nucleic Acids Res. 2019;47:D801–6.

97. Febbo PG, Mulligan MG, Slonina DA, Stegmeir K, Di Vizio D, Martinez PR, et al. Literature Lab: a method of automated literature interrogation to infer biology from microarray analysis. BMC Genomics. 2007;8:461.

98. Shannon P, Markiel A, Ozier O, Baliga NS, Wang JT, Ramage D, et al. Cytoscape: A Software Environment for Integrated Models of Biomolecular Interaction Networks. Genome Res. 2003;13:2498–504.

