## Additional File 1 for "Differential regulation of the immune system in a brain-liver-fats organ network during short term fasting"

[1] Sidra Medicine, PO Box 26999, Doha, Qatar.

[2] Institute of Epigenetics and Stem Cells, Helmholtz Zentrum München, Marchioninistraße 25, 81377 München, Germany

[3] Institute of Functional Epigenetics, Helmholtz Zentrum München, Ingolstädter Landstraße 1, 85764 Neuherberg, Germany

[4] Institute of Computational Biology, Helmholtz Zentrum München, Ingolstädter Landstraße 1, 85764 Neuherberg, Germany

[5] Wellcome Sanger Institute, Wellcome Genome Campus, Hinxton, Cambridge, CB10 1SD, UK.

[6] Monell Chemical Senses Center, 3500 Market Street, Philadelphia, PA 19104, USA.

[#] equal contribution (co-first)

† equal contribution (co-senior)

**Additional file 1: Supplementary Figures S1-S6**

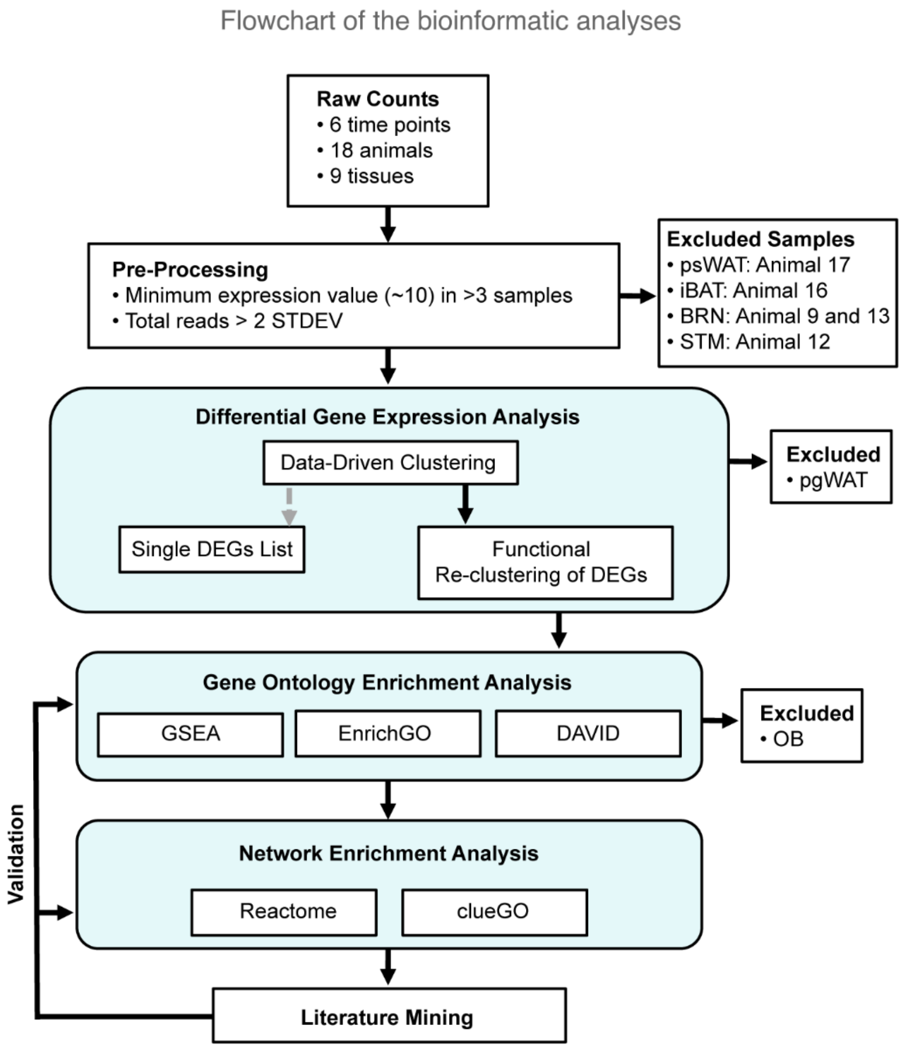


**Additional file 1: Figure S1. Schematic of the bioinformatic workflow.**

Genes and samples were filtered for minimum expression threshold, replicate numbers, and outlier exclusion. A data-driven approach was then used to group samples into Phases to derive the differentially expressed genes which were then functionally clustered by their semantic similarities prior to gene ontology analysis. A group of four highly overlapping organs (BRN, LIV, iBAT and psWAT) was carried forward to protein enrichment analyses and the enrich processes validated using known literature.

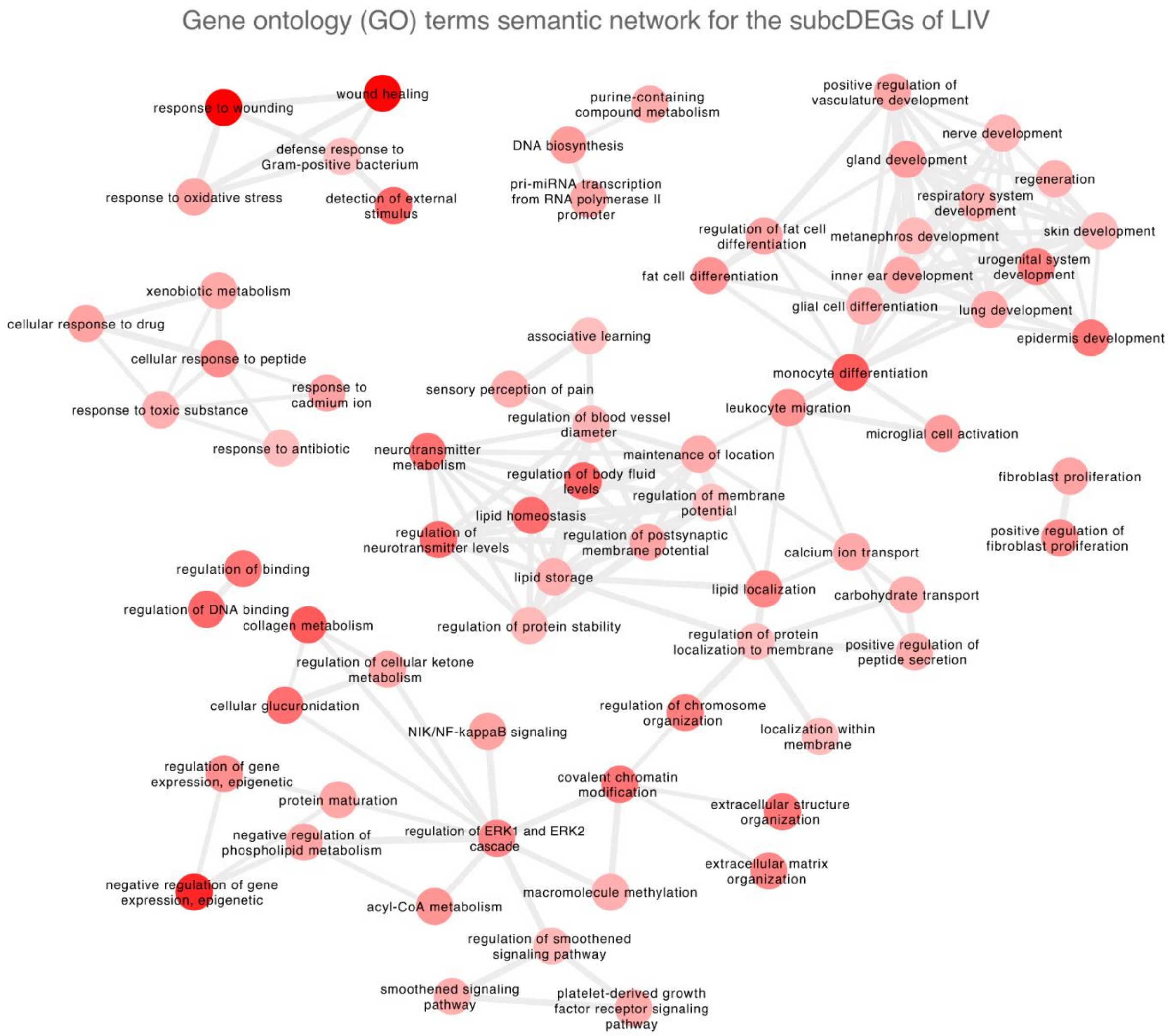


**Additional file 1: Figure S2. Semantic network of the gene ontology (GO) terms enriched in the liver.** Nodes are GO terms and the intensity of the colour indicates enrichment (-$\log_{10}$ p-value, darker shade indicates lower p-values). Highly similar GO terms are linked by edges, where the line width indicates the degree of similarity.

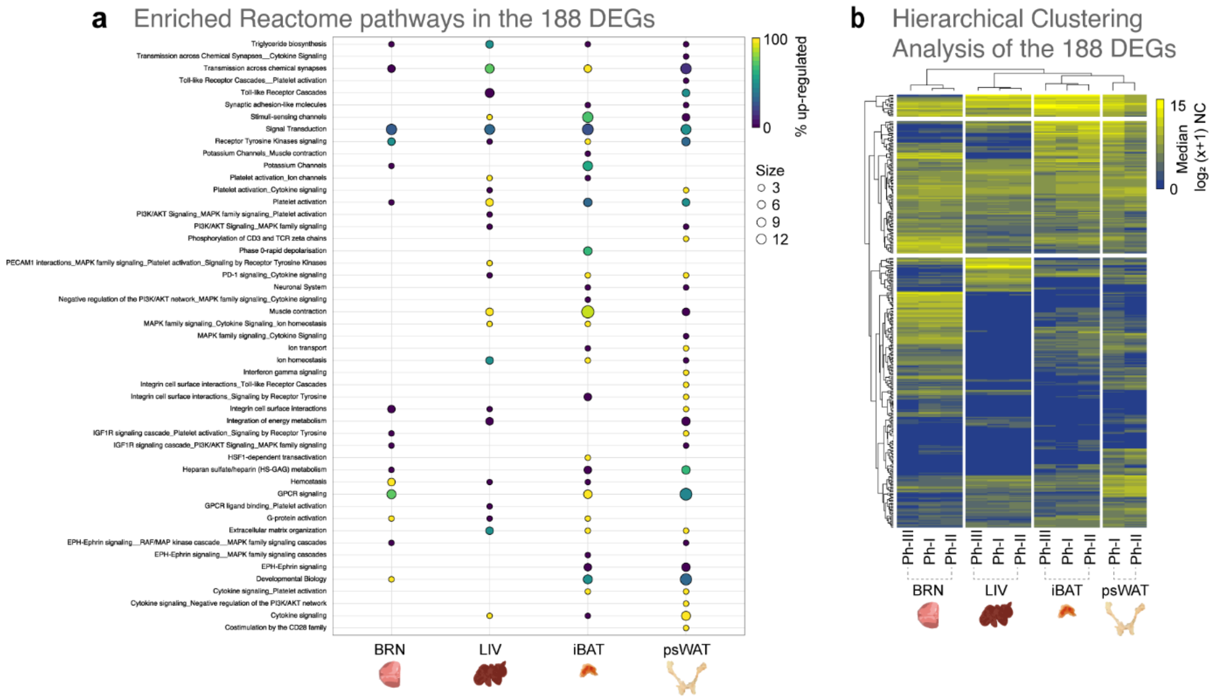


**Additional file 1: Figure S3. Gene expression of the *brain-liver-fats* organ network.**

**a)** Percent of genes up-regulated in the summarized enriched Reactome terms of the *brain-liver-fats* organ network (Fig. 4a) in BRN, LIV, iBAT, and psWAT. Similar/redundant Reactome terms were combined for simplicity of representation.

**b)** Hierarchical clustering analysis of the 188 genes from the *brain-liver-fats* organ network. Median mRNA expression levels are represented on a $\log_2$ (x+1) scale of normalized counts (NC) (0 - not expressed; 15 - highly expressed) per fasting Phases (Ph) and across the four organs.

**a** LitLab™ Table for the *brain-liver-fats* organ network

| Physiology-MeSH | Association | Gene X Term Abstracts | Term Abstracts | Nonzero Genes | Score | P-Value |
|---|---|---|---|---|---|---|
| Nervous System Physiological Phenomena | Strong | 23655 | 728532 | 172 | 3.25 | 0.0006 |
| Drug Tolerance | Strong | 705 | 11987 | 60 | 2.94 | 0.0017 |
| Pain | Strong | 15864 | 664201 | 156 | 2.69 | 0.0035 |
| Electrophysiological Phenomena | Strong | 11468 | 270347 | 159 | 2.63 | 0.0042 |
| Pain Threshold | Strong | 1055 | 14698 | 72 | 2.58 | 0.0049 |

| Pathway-MeSH | Association | Gene X Term Abstracts | Term Abstracts | Nonzero Genes | Score | P-Value |
|---|---|---|---|---|---|---|
| Calcium Signaling | Strong | 24928 | 182977 | 177 | 3.02 | 0.0012 |
| BDNF | Strong | 12902 | 12913 | 113 | 2.75 | 0.003 |
| Nifedipine | Strong | 506 | 5336 | 63 | 2.57 | 0.0051 |
| Alpha-Adrenergic Signaling | Strong | 122 | 1707 | 36 | 2.51 | 0.006 |
| Butirosin/Neomycin Biosynthesis | Moderate | 229 | 3408 | 62 | 2.98 | 0.0014 |
| Trka | Moderate | 3222 | 3222 | 62 | 2.57 | 0.005 |
| Cardiac Calcium Regulation | Moderate | 3434 | 21350 | 122 | 2.49 | 0.0063 |
| EPHA | Moderate | 1117 | 2659 | 65 | 2.4 | 0.0081 |
| Cardiac Muscle Contraction | Moderate | 1575 | 13542 | 98 | 2.36 | 0.0091 |
| Beta-Agonist/Beta-Blocker | Moderate | 225 | 1860 | 49 | 2.36 | 0.0093 |
| Fatty Acid Biosynthesis | Moderate | 1378 | 5610 | 48 | 2.3 | 0.0109 |
| Nerve Growth Factor | Moderate | 18204 | 38144 | 158 | 2.29 | 0.011 |
| Mitochondrial Carnitine Palmitoyltransferase | Moderate | 761 | 3663 | 64 | 2.29 | 0.0111 |
| Ca - Calmodulin | Moderate | 853 | 3969 | 94 | 2.24 | 0.0124 |
| T Cell CD3 | Moderate | 3885 | 9572 | 74 | 2.2 | 0.0138 |
| Palmitate biosynthesis | Moderate | 900 | 7196 | 63 | 2.17 | 0.015 |
| SRC | Moderate | 195 | 394 | 29 | 2.16 | 0.0155 |
| Drug Addiction | Moderate | 142 | 865 | 35 | 2.16 | 0.0155 |
| Toll-Like Receptor Pathway | Moderate | 20621 | 35256 | 118 | 2.11 | 0.0176 |
| Antigen | Moderate | 64961 | 344214 | 179 | 2.08 | 0.0189 |
| T-Cell Receptor and CD3 Complex | Moderate | 2500 | 6349 | 54 | 2.06 | 0.0195 |
| Lymphocyte Diapedesis | Moderate | 5427 | 13440 | 84 | 2.05 | 0.0204 |
| Lymphocyte Adhesion | Moderate | 5415 | 13368 | 83 | 2.04 | 0.0204 |
| TCR Signaling | Moderate | 4205 | 12513 | 82 | 2.04 | 0.0209 |

**b** Major categories enriched

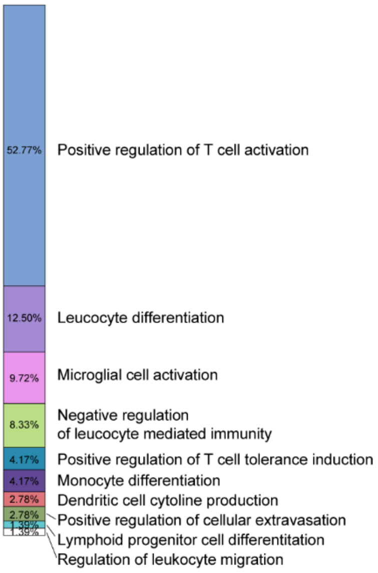


**c** Gene ontology (GO) terms Immune upregulated

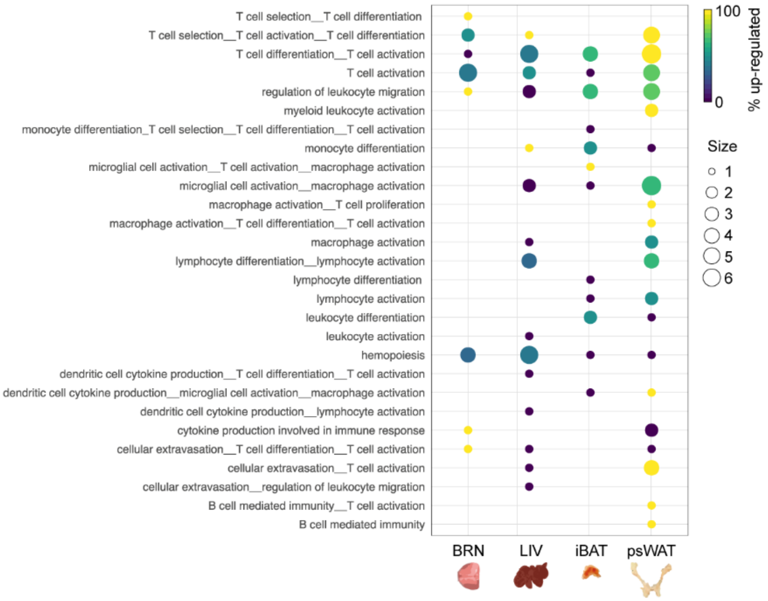


**Additional file 1: Figure S4. Short-term fasting modulates immune processes in the *brain-liver-fast* organ network.**

**a)** Summary of the significant Physiology and Pathway-specific Medical Subject Headings (MeSHs) associated with the 188 organ network genes resulting from the association analysis performed using LitLab™.

**b)** The representative categories and percentages of the immune-specific GO terms enriched from the 96 genes extracted from the 37 overlapping GO terms from BRN, LIV, iBAT and psWAT.

**c)** Proportion of the up-regulated gene from the 96 genes from the organ network that resulted in a significant immune-specific GO network enrichment.

| Tissue | ImmuneGene | Comparison | $log_2FC$ | padj | Function |
|---|---|---|---|---|---|
| BRN | Tff2 | III vs I | 4.06 | 0.02 | Inhibits gastric acid secretion and is expressed in gastrointestinal mucosa; negative regulation of macrophage activation; regulates a function/process that happens in peritoneal cavity (J:116628). |
| | Dmbt1 | III vs I | 3.66 | 0.00 | Receptor that bind surfactant protein D, may play a role in cell-cell interaction; contributes to protection from infection via its presence in mucosal fluid (28489917); endothelium-derived ECM protein; KO associated with increase susceptibility to inflammation and increase TNF, IL6, NOD2 expression during inflammation; antimicrobial humoral immune response mediated by antimicrobial peptide; induction of bacterial agglutination (J:164563). |
| | Ctla2a | III vs I | 3.21 | 0.00 | Immunosuppressive, promote T reg and suppress T effector via induction of TGF-beta secretion (19801522); distributed in mouse brain (18342295).; Maintain normal growth and development (27586811); regulation of regulatory T cell differentiation (J:171531). |
| | Gng13 | III vs I | 2.87 | 0.00 | Essential for olfactory function in mice (23637188, 30157062); taste transduction cascade may be involved in spermatogenesis (22266327). |
| | Reg1 | III vs I | 2.86 | 0.02 | Immunomodulator of IL36 signal (29339122); involved in beta-cell regeneration pathway (23986224, 29935186); antimicrobial humoral immune response mediated by antimicrobial peptide (J:265628). |
| LIV | Myc | II vs I | 2.77 | 0.00 | TF. Cell cycle/apoptosis. Evidence for non-AUG translated protein isoform under stress and nutrient deprivation with unspecified molecular mechanism; negative regulation of monocyte differentiation (J:164563). |
| | Egr1 | III vs II | 2.64 | 0.00 | TF. Early growth factor-1. T cell differentiation (J:78245). |
| | Gadd45g | III vs I | 2.07 | 0.01 | Growth arrest 45g. Downregulation of Gadd45g contributes to JAK/STAT3 activation and cellular senescence avoidance (23897841). Gadd45g acts as a cold-induced activator of BAT thermogenesis via MAPK/p38 pathway (25071184); T-helper 1 cell differentiation (J:79231). |
| | Lpin1 | III vs II | 1.69 | 0.00 | Lipin. Down-regulation of hepatic Lipin1 expression leads to less adiposity as well as a decrease in TG level in the liver and blood circulation (27725442). |
| | Gadd45b | III vs II | 1.54 | 0.00 | Growth arrest 45b. Fasting-induced liver GADD45β restrains hepatic fatty acid uptake and improves metabolic health (27137487). |
| iBAT | Fzd9 | III vs I | 4.23 | 0.00 | Neurone regeneration, neural precursor cells, cortex/spiral ganglion neurons (19716861, 15354291, 11685582); B cell differentiation in the bone marrow (J:98133). |
| | Cd8a | III vs I | 3.65 | 0.01 | T cell receptor; adaptive immune response, cytotoxic T cell differentiation, T cell mediated immunity (J:60000, J:265628, J:68956, J:143981, J:155856). |
| | Mb | III vs II | 3.40 | 0.00 | Myoglobin, protect cardiac fonction in heypoxia and disease and induce FA metabolism via b-oxidation (28230173); contributes to vasodilation via nitrites (20889759); enucleate erythrocyte differentiation (J:95767). |
| | Wnt4 | III vs I | 3.34 | 0.00 | Expression in DC mediate suppression of Th2 immunity (31175162); prevents inflammation by inhibiting NFkB (25108526). |
| | Thbs4 | III vs I | 3.24 | 0.00 | Thrombosopondin-4, proangiogenic, recruits macrophoge and help restenosis; plays a role in macrophage migration in adipose tissue (26868511); positive regulation of neutrophil chemotaxis (J:164563). |
| psWAT | Mmp8 | II vs I | 5.41 | 0.00 | ECM cleavage and promotes macrophage differentiation to M2 via increased in TGFb bioavailability (26092731); positive regulation of microglial cell activation and neuroinflammatory response; is a part of cellular response to lipopolysaccharide (J:265613). |
| | Retnlg | II vs I | 5.37 | 0.00 | Elevated in high-fat-fed/obeses mice in intestinal tract and bone marrows, potential regulation of insulin sensitivity (15834545); may play a role in promyelocytic differentiation (163736900); myeloid dendritic cell chemotaxis (J:89564). |
| | S100a8 | II vs I | 4.16 | 0.00 | Modulate neuroinflammation in sepsis survivor mice, including granulocyte recruitment and priming of microglial-reactive oxygen species and cytokine production (29563178); reduce acute lung injuries (27365293). |
| | S100a9 | II vs I | 4.05 | 0.00 | Necessary in the recruitment of neutrophils and for priming production of reactive oxygen species and TNF-α secretion in microglia and macrophages (29563178). Promotes MDSC expansion and immunosuppresion in late/chornic sepsis (31078118). |
| | Tlr9 | II vs I | 3.35 | 0.00 | activation of innate immune response (J:240584); defense response to virus (J:265628 , J:131989). |

**Additional file 1: Figure S5. Immune-related DEGs in BRN, LIV, iBAT and psWAT.**

Top 5 up-regulated immune-related genes among DEGs of the four organs (BRN, LIV, iBAT and psWAT) and their literature-supported functions. The immune-related functions of each gene were extracted from NCBI and MGI databases, as indicated by their PubMed IDs and reference IDs.

| Tissue | ImmuneGene | Comparison | $log_2FC$ | padj | Function |
|---|---|---|---|---|---|
| BRN | Gh | III vs I | -4.69 | 0.04 | Growth hormone |
| | Ly96 | III vsII | -4.12 | 0.04 | Promotes inflammation via tlr4-nox4 pathways (29111459); promotes inflammation; toll-like receptor 4 signaling pathway (J:265628, J:222669). |
| | Fgf18 | III vs I | -3.34 | 0.01 | Fibroblast growth factor; growth factor activity. |
| | Igf2 | III vs I | -3.10 | 0.03 | Insulin growth factor, involved in development and growth; negative regulation of natural killer cell mediated cytotoxicity (J:155856) ; positive regulation of activated T cell proliferation (J:164563). |
| | Ifi203 | III vs I | -3.09 | 0.03 | Neural development (30635555); activation of innate immune response (J:72247). |
| LIV | Dmbt1 | II vs I | -5.93 | 0.01 | Receptor that bind surfactant protein D, may play a role in cell-cell interaction; contributes to protection from infection via its presence in mucosal fluid (28489917); endothelium-derived ECM protein; KO associated with increase susceptibility to inflammation and increase TNF, IL6, NOD2 expression during inflammation.antimicrobial humoral immune response mediated by antimicrobial peptide; induction of bacterial agglutination (J:164563). |
| | Lcn2 | III vs I | -3.07 | 0.02 | Innate immune protein induce by ROS (30279516). |
| | Orm2 | III vs I | -2.42 | 0.04 | Transcription of Orm2 induced during acute phase; regulation of immune responses (J:72247) |
| | Chrna4 | III vs I | -2.32 | 0.02 | Cholinergic receptor, nicotinic, alpha polypeptide 4; B cell activation,has the participant spleen (J:103908). |
| | Prtn3 | III vs I | -2.11 | 0.04 | Proteinase 3. Similarly to caspase-1, PR3 can process IL-1b and IL-18 to active forms, promoting inflammation. Associated with insulin resistance. KO is associated with reduced lipids in liver after high fat diet (27261776); mature conventional dendritic cell differentiation, neutrophil extravasation (J:164563). |
| iBAT | Alpk1 | III vs I | -5.08 | 0.00 | Alpha-kinase 1 is a cytosolic innate receptor for bacterial saccharides; suppresses inflammation via downregulation of IL12/Th1 axis (30228258); mediate inflammation via NKFB (30111836). |
| | Chst3 | III vs I | -4.74 | 0.00 | Carbohydrate sulfotransferase 3 diminishes macrophage accumulation via MMP-9 downregulation (26646821); T cell homeostasis, happens in spleen,affects a population of T cell (J:73698). |
| | Aim2 | III vs I | -4.41 | 0.00 | ds-DNA sensing molecule that mediates inflammasome activation; induction leads to IL-1b production, pyroptosis, and bacterial infection resistance; activation of innate immune response (J:265628, J:147175, J:164563) |
| | H60b | III vs I | -4.30 | 0.00 | Histocompatibility 60b, ligand for NKG2D on NK cells rendering those cells targetable (18209064); natural killer cell mediated cytotoxicity (J:131505). |
| | Ide | III vs I | -4.24 | 0.00 | Insulin degrading enzyme. Lower levels of IDE contributes to favorable blood glucose dynamics during protein-restricted mice (25036874); antigen processing and presentation of endogenous peptide antigen via MHC class I (J:164563). |
| psWAT | Trim29 | II vs I | -4.72 | 0.00 | Negative reguation of type 1 IFN in response to RNA Virus(29769269); inhibits innate immune response in DNA virus infections (29038422) |
| | Krt75 | II vs I | -4.65 | 0.00 | Keratin, related to hair follicles; hematopoietic progenitor cell differentiation (J:202604). |
| | Btn1a1 | II vs I | -4.32 | 0.04 | Expressed in mammary gland and interact with milk fat globule; regulation of milk-lipid droplet (15226505); inhibitor of T cell activation (20208008); negative regulation of activated T cell proliferation (J:160063). |
| | Bpifa1 | II vs I | -4.12 | 0.01 | Regulates neutrophil recruitment and interferon signaling during inflammation; defeciency causes susceptibility to infection; antibacterial humoral response (J:195535, J:164563). |
| | Otop1 | II vs I | -3.72 | 0.00 | Otopetrin, component of a counterinflammatory pathway that is induced in WAT during obesity; elevated in obese mice and induced by TNFa in adipocytes (24379350); pPhysically interact with STAT1 and attenuates obesity-induced adipose tissue inflammation; negative regulation of interferon-gamma-mediated signaling pathway (J:229138). |

**Additional file 1: Figure S6. Immune-related DEGs in BRN, LIV, iBAT and psWAT.**

Top 5 down-regulated immune-related genes among DEGs of the four organs (BRN, LIV, iBAT and psWAT) and their literature-supported functions. The immune-related functions of each gene were extracted from NCBI and MGI databases, as indicated by their PubMed IDs and reference IDs.
